# The NAD^+^-dependent deacetylase Sir2 enables evolution of new traits by regulating distinct gene sets in two yeast species, *Saccharomyces cerevisiae* and *Kluyveromyces lactis*

**DOI:** 10.1101/734608

**Authors:** Kristen M. Humphrey, Lisha Zhu, Meleah A. Hickman, Shirin Hasan, Haniam Maria, Tao Liu, Laura N. Rusche

**Affiliations:** Department of Biological Sciences, University at Buffalo State University of New York, Buffalo, NY, 14260; Department of Biochemistry, University at Buffalo State University of New York, Buffalo, NY, 14203; Department of Biochemistry, Duke University, Durham, NC, 27710

**Author notes:** Department of Cancer Genetics & Genomics, Roswell Park Cancer Institute, Buffalo, NY, 14203. School of Biomedical Informatics, University of Texas Health Science Center at Houston, Houston, TX, 77030. Department of Biology, Emory University, Atlanta, GA, 30322. Department of Medicine, Northwestern University School of Medicine, Chicago, IL, 60611. Department of Biostatics & Bioinformatics, Roswell Park Cancer Institute, Buffalo, NY, 14203. Data available in public repositories: GSE85574, GSE84552, GSE92930, GSE86149, and GSE84403. Corresponding author: Laura Rusche, 109 Cooke Hall, Buffalo. NY 14260.

**Keywords:** Sirtuin, Sum1, nicotinamide adenine dinucleotide, sporulation

## Abstract

Evolutionary adaptation increases the fitness of an organism in its environment. It can occur through rewiring of gene regulatory networks, such that an organism responds appropriately to environmental changes. We investigated whether sirtuin deacetylases, which repress transcription and require NAD^+^ for activity, could facilitate the evolution of potentially adaptive responses by serving as transcriptional rewiring points. If so, bringing genes under the control of sirtuins could enable organisms to mount appropriate responses to stresses that decrease NAD^+^ levels. To explore how the genomic targets of sirtuins shift over evolutionary time, we compared two yeast species, *Saccharomyces cerevisiae* and *Kluyveromyces lactis* that display differences in cellular metabolism and lifecycle timing in response to nutrient availability. We identified sirtuin-regulated genes through a combination of chromatin immunoprecipitation and RNA expression. In both species, regulated genes were associated with NAD^+^ homeostasis, mating, and sporulation, but the specific genes differed. In addition, regulated genes in *K. lactis* were associated with other processes, including utilization of non-glucose carbon sources, heavy metal efflux, DNA synthesis, and production of the siderophore pulcherrimin. Consistent with the species-restricted regulation of these genes, sirtuin deletion impacted relevant phenotypes in *K. lactis* but not *S. cerevisiae*. Finally, sirtuin-regulated gene sets were depleted for broadly-conserved genes, consistent with sirtuins regulating processes restricted to a few species. Taken together, these results are consistent with the notion that sirtuins serve as rewiring points that allow species to evolve distinct responses to low NAD^+^ stress.

## INTRODUCTION

Evolutionary adaptation is the process by which species acquire traits that make them better suited to a particular environment. At the molecular level, adaptation can involve the acquisition of new protein functions or new gene expression patterns (OHNO 1970; KING AND WILSON 1975). In the case of new gene expression patterns, particular transcriptional regulators might be frequent rewiring points for such changes. For example, a regulator that responds to a particular stress could be redirected to new genes to invoke a distinct biological response to that stress. In this study, we explored the possibility that sirtuin deacetylases are such regulators.

Sirtuins are NAD^+^-dependent deacetylases that have been identified in all kingdoms of life (GREISS AND GARTNER 2009). These enzymes have two characteristics that are consistent with being rewiring points. First, sirtuins require NAD^+^ for activity (IMAI *et al*. 2000) and are therefore proposed to respond to stresses that impact intracellular NAD^+^ levels. In particular, because NAD^+^ is a key redox carrier in central metabolism, changes in metabolic flux due to nutrient perturbations could impact NAD^+^ availability and hence the deacetylase activity of sirtuins. At this time, the conditions that cause low intracellular NAD^+^ in yeast are not well understood. However, there is evidence that intracellular NAD^+^ levels impact Sir2 activity. For example, genetic perturbations of the enzymes and transporters that maintain intracellular NAD^+^ alter Sir2 activity (SMITH *et al*. 2000; ANDERSON *et al*. 2002; BELENKY *et al*. 2007), as does the presence or absence of NAD^+^ precursors in the growth medium (BELENKY *et al*. 2007). Moreover, intracellular NAD^+^ is increased in the absence of inositol (LEE *et al*. 2013) and in aging cells (ASHRAFI *et al*. 2000) and these increases correspond with enhanced Sir2 activity.

A second characteristic of sirtuins consistent with being rewiring points is that they have diverged to deacetylate a wide range of substrates. Some sirtuins, including those examined here, deacetylate histones and are targeted to specific genomic locations where they repress transcription. Thus, these sirtuins are proposed to connect gene expression to the metabolic state of the cell by sensing NAD^+^ levels. Therefore, species could evolve distinct biological responses to stresses that perturb NAD^+^ levels by bringing new genes under the control of sirtuins. Consistent with this idea, we have observed that in the *Candida* clade of yeast the genomic targets of sirtuins vary from one species to another (FROYD *et al*. 2013; KAPOOR *et al*. 2015; RUPERT *et al*. 2016).

To explore how the genes targeted by sirtuins have shifted over evolutionary time, we compared two genetically tractable yeast species, *Saccharomyces cerevisiae* and *Kluyveromyces lactis*. *S. cerevisiae* Sir2 (ScSir2) deacetylates histone tails (BRAUNSTEIN *et al*. 1993; IMAI *et al*. 2000) and, as part of a complex with Sir3 and Sir4, forms transcriptionally repressive chromatin at telomeres and the cryptic mating-type loci (RINE AND HERSKOWITZ 1987; GOTTSCHLING *et al*. 1990). ScSir2 also acts at the tandem rDNA repeats where it reduces recombination (GOTTLIEB AND ESPOSITO 1989). A paralog of Sir2, Hst1 (<u>H</u>omolog of <u>S</u>ir <u>T</u>wo), arose in a whole genome duplication (BYRNE AND WOLFE 2005). ScHst1 acts with the DNA-binding protein Sum1 to repress mid-sporulation and NAD^+^ biosynthetic genes (XIE *et al*. 1999; BEDALOV *et al*. 2003). We compared the targets of ScSir2 and ScHst1 in *S. cerevisiae* with those of KlSir2 in *K. lactis.* This species did not undergo the duplication that led to Sir2 and Hst1. Like ScSir2, KlSir2 acts at the telomeres, cryptic mating-type loci, and rDNA; and like ScHst1, KlSir2 acts at mid-sporulation genes (ASTROM *et al*. 2000; HICKMAN AND RUSCHE 2009).

An unresolved question is whether the sets of genes repressed by Sir2 and Hst1 in *S. cerevisiae* and *K. lactis* differ functionally, enabling each species to respond in a distinct way to low intracellular NAD^+^. There are several key differences between these species that are connected to cellular metabolism and nutrient availability. For example, in the presence of oxygen, *S. cerevisiae* processes sugars through fermentation whereas *K. lactis* favors respiration (KIERS *et al*. 1998). These distinct metabolic strategies might require different responses to low NAD^+^ levels. A second difference is the coordination of the sexual cycle with nutrient availability. Newly germinated *S. cerevisiae* spores, which are haploid, mate readily in rich nutrients. The resulting diploid cells propagate mitotically until nutrients become scarce, at which point they undergo meiosis and sporulation. In contrast, newly germinated *K. lactis* spores do not mate readily but instead propagate in the haploid state until nutrients become scarce (HERMAN AND ROMAN 1966; BOOTH *et al*. 2010). They then mate and proceed directly to meiosis and sporulation.

To determine whether Sir2 and Hst1 regulate distinct genes that would be advantageous for the different metabolic and life cycle strategies of *S. cerevisiae* and *K. lactis*, we defined the gene sets regulated by Sir2 and Hst1 in both species. To do so, we used a combination of chromatin immunoprecipitation and RNA expression analyses. Regulated genes in both species are involved in NAD^+^ homeostasis, mating, and sporulation. However, the specific genes that are regulated differ. In addition, in *K. lactis*, regulated genes are associated with processes not regulated in *S. cerevisiae*, including utilization of non-glucose carbon sources, heavy metal efflux, DNA synthesis, and production of the siderophore pulcherrimin. Consistent with the species-restricted regulation of these genes, sirtuin deletion impacted relevant phenotypes in *K. lactis* but not *S. cerevisiae*. We also found that the gene sets regulated by Sir2 and Hst1 are depleted for widely-conserved genes. Taken together, these results are consistent with the notion that sirtuins can serve as rewiring points that allow species to evolve distinct responses to low NAD^+^ stress.

## MATERIALS AND METHODS

### Plasmids and yeast strains

Plasmids used in this study are listed in Table S1. The details of construction are provided in the supplemental materials and methods. Yeast used in this study are listed in Table S2. Most *K. lactis* strains were derived from Os334 and Os335 (HEINISCH *et al*. 2010), which are congenic with the type strain CBS2359. *S. cerevisiae* strains were derived from the laboratory strain W303-1b. The details of strain construction are provided in the supplemental materials and methods.

### Yeast growth and transformation

Yeast were grown at 30° in YPD (1% yeast extract, 2% peptone, 2% glucose) unless otherwise stated. *S. cerevisiae* cells were transformed using the PEG-LiOAc method (SCHIESTL AND GIETZ 1989), and *K. lactis* cells were transformed by electroporation (HICKMAN AND RUSCHE 2009).

To follow growth in various carbon sources, cultures were grown in YM (0.67% yeast nitrogen base without amino acids) with the desired carbon source (2% glucose or 3% glycerol). For *S. cerevisiae*, YM was supplemented with histidine, leucine, lysine, and tryptophan. Overnight (glucose) or three-day (glycerol) cultures were diluted 50% in the same medium and grown an additional 3 hours at 30°. Cells were then diluted to OD 0.05 and placed in a 96-well clear-bottom plate containing YM supplemented with the desired carbon source. The OD_600_ of each culture was recorded at consistent time intervals at 30° on a SpectraMax i3x microplate reader using the SoftMaxPro 6.5.1 software. To follow growth in HU or sodium arsenate, cells were grown overnight in YPD, diluted to OD = 0.3, and allowed to grow 4 hours at 30°. Cells were then diluted to OD = 0.05 in different concentrations of HU or sodium arsenate, and the growth was recorded as described above. To assess pulcherrimin production, cells grown overnight in YPD, diluted to OD_600_ = 0.3 in YPD, and grown four hours. Cells were collected, washed, and re-suspended in PBS at 1 OD/ml. 10μL of cells were spotted onto YPD plates supplemented with 3.7 mM FeCl_3_·6H_2_O. The plates were incubated two days and imaged.

### RNA isolation and sequencing

For sequencing, RNA was isolated from *S. cerevisiae* strains LRY3093 and 3099 and *K. lactis* strains LRY2835, 2849, and 2850 (2012 data set) or LRY2835, 2850, 3096, and 3098 (2016 data set) as previously described (SCHMITT *et al*. 1990). Cells were grown in YPD and harvested in mid-log phase at OD_600_ about 1. DNA was removed from 10 μg RNA using Turbo DNase (Life Technologies), according to the manufacturer’s instructions. One set of *K. lactis* RNA samples was processed for sequencing using the TruSeq non-stranded RNA Library Prep Kit (Illumina), and 50 bp single end sequencing was performed on an Illumina HiSeq2000 machine at the Duke IGSP sequencing facility. A second set of *K. lactis* samples and the *S. cerevisiae* samples were processed at the University at Buffalo Genomics and Sequencing Core facility and sequenced on an Illumina HiSeq2500.

### Chromatin IP and processing for microarray or sequencing

For the ChIP-Seq experiment from *S. cerevisiae*, ScSir2-HA was immunoprecipitated from LRY1926, ScHst1-HA was immunoprecipitated from LRY558, and LRY1009 was used for mock IP and input DNA. Chromatin IP and processing is described in the supplemental methods. Library preparation and sample barcoding was done at the Next-Generation Sequencing facility at University at Buffalo. The samples were then sequenced on an Illumina HiSeq2500 using 50 bp single-end sequencing.

For the ChIP on Chip experiment, KlSir2-HA was immunoprecipitated from strains LRY2021 or LRY2022. For LRY2021, the IP sample was labeled with Cy5 and the input was labeled with Cy3. For LRY2022, the dyes were swapped. Labeled DNA was hybridized to a custom ChIP-on-Chip 2×105K microarray (Agilent G4498A), designed with 102,839 60-nucleotide probes tiled across the *K. lactis* genome spaced approximately every 100 bp (AMADID 018357). Chromatin IP was conducted as previously described (HICKMAN AND RUSCHE 2009) and processed as described in the supplemental methods.

### Bioinformatic analysis

ChIP-Seq reads of ScSir2 and ScHst1 were mapped to the *S. cerevisiae* reference genome downloaded from SGD database (http://www.yeastgenome.org/strain/S288C/overview) using BWA v0.7.7-r441 (LI AND DURBIN 2009). For calling enriched peak regions, MACS2 v 2.0.10 (ZHANG *et al*. 2008) was used with genomic input as control, and the parameters used were “-B --nomodel --extsize 200 -q 0.01 -g 12157105”. This analysis identified 159 ScSir2 peaks and 692 ScHst1 peaks. Genes associated with these peaks were defined as those for which the gene body intersected with a peak in addition to those for which the start codon was within 1 kbp of a KlSir2 peak.

ChIP-on-chip signals were mapped to the *K. lactis* reference genome (https://www.ncbi.nlm.nih.gov/genome/?term=txid28985%5Borgn%5D), and KlSir2 binding sites were called using MA2C software (SONG *et al*. 2007). The normalization method was set as “Robust,” the FDR (False Discovery Rate) cutoff for ChIP-enriched regions was 0.05, the BANDWIDTH was set to 500 bp, the number of MIN_PROBES was 5, and the MAX_GAP was set as 250 bp. This analysis identified 460 KlSir2 peaks. Genes associated with these peaks were defined using the same criteria as for *S. cerevisiae*.

For RNA-Seq, the raw reads from single-end sequencing were mapped to the *K. lactis* and *S. cerevisiae* reference genomes, allowing no more than 2 mismatches, using tophat v2.0.10 (KIM *et al*. 2013). FPKM (Fragments Per Kb per Million reads) were obtained using cufflinks version 2.2.1 (TRAPNELL *et al*. 2013) and count data were obtained using HTSeq-count (ANDERS *et al*. 2015) with alignment quality cutoff set to 10. The gene annotation file for *K. lactis* was obtained from Genolevures database (http://genolevures.org/index.html<u>#</u>) and for *S. cerevisiae* was obtained from Ensembl database (http://ftp.ensembl.org/pub/release-66/gtf/saccharomyces_cerevisiae/). edgeR (ROBINSON *et al*. 2010) was applied to detect genes differentially expressed between wild-type and knockout strains, with FDR cutoff of 0.05 and the absolute fold-change larger than or equal to 2.

To identify *S. cerevisiae* orthologs of *K. lactis* genes, each *K. lactis* gene served as a BLASTP query, and the top *S. cerevisiae* hits (≥70% of the maximum score for that search) were identified. Next, these *S. cerevisiae* genes were used as queries, and the top *K. lactis* hits were identified. If the second BLASTP search returned to the starting *K. lactis* gene, the genes from the two species were concluded to be orthologs. In some cases, the BLASTP search identified a multi-gene family or two paralogs that arose through duplication in the *S. cerevisiae* lineage. These genes were all taken as orthologs. Most of the ortholog assignments were consistent with the Yeast Gene Order Browser (BYRNE AND WOLFE 2005), which considers gene order as well as homology. After manual refinement, including the use of gene order to assign orthologs to some short, rapidly evolving genes, we identified *S. cerevisiae* orthologs for 109 of the 175 KlSir2-regulated genes (Table S7).

### Mating assays

For *S. cerevisiae* mating assays, cells of both mating types were grown separately overnight in YPD at 30° with shaking. Cultures were diluted by 50% in YPD and incubated for 3 hours, spun down, and resuspended at 10 OD/mL in YPD. A lawn was prepared using 200μL (2 OD of cells) of one mating-type spread on YM plates supplemented with histidine, tryptophan, and leucine. The other stain was then used to make five 10-fold serial dilutions. 200μL of the most dilute culture (2×10^-5^ OD) was spread on YPD plates to determine the number of viable cells and over the lawn to determine the number of cells that could mate. Three mating lawns and one YPD plates were counted for each mating combination.

For *K. lactis* mating assays, *MAT**a*** cells expressed mCherry, and *MATα* cells expressed yEGFP. Cells of both mating types were co-cultured on malt extract (5% malt extract and 3% agar) for three days at 30°. Cells were resuspended in 500 μL of sterile distilled water and then sonicated using a soniprep 150 sonicator for 5 seconds at an amplitude of 5 microns. Finally, cells were examined using ImageStream cytometry (ImageStream Mark II Imaging flow cytometer, EMD Millipore) with the following settings: 488nm laser at 100 watts, 561nm laser at 150 watts, 60X amplification. Images were collected for brightfield in channels 1 and 9, side scatter in channel 6, yEGFP in channel 2, and mCherry in channel 4. 100,000 events were collected for each sample. IDEAS software (EMD Millipore) was used to identify cells which were in focus and showed both red and green fluorescence. Each of these events were manually examined for an hourglass appearance typical of zygotes.

### Sporulation assay

Diploid cells were freshly grown from frozen glycerol stocks. After overnight growth on YPD plates, cells were patched onto KOAc plates (1% KOAc and 3% agar) and incubated at 30°. At two hour intervals, cells were resuspended in sterile distilled water, vortexed, and examined under a light microscope. All cells in three fields of vision were scored as either having sporulated (tetrad morphology) or not. The experiment was conducted twice with four biological replicates (diploid strains) each time.

### Data and reagent availability statement

Strains and plasmids are available upon request. Gene expression data are available at GEO with the accession numbers: GSE92930, GSE86149, and GSE84403. ChIP-Seq data are available at GEO with the accession number GSE84552. ChIP-on-Chip data are available at GEO with the accession number: GSE85574.

Table S1 lists plasmids used in this study. Table S2 lists yeast strains used in this study. Table S3 contains RNA-Seq and ChIP-Seq data for all annotated *S. cerevisiae* genes. Table S4 contains RNA-Seq and ChIP-Chip data for all annotated *K. lactis* genes. Table S5 contains descriptions and comparative information for ScSir2-regulated genes. Table S6 contains descriptions and comparative information for ScHst1-regulated genes. Table S7 contains descriptions and comparative information for KlSir2-regulated genes identified based on 2016 RNA-Seq data. Table S8 contains descriptions and comparative information for KlSir2-regulated genes identified based on 2012 RNA-Seq data. Table S9 is the basis for figure 3. It includes RNA-Seq and ChIP data for the genes known to act in each pathway represented in the figure.

## RESULTS

### Identification of genes regulated by ScSir2, ScHst1, and KlSir2

To identify the genes regulated by Sir2 and Hst1 in *S. cerevisiae* and *K. lactis*, we combined data from two experiments for each species. First, we mapped genomic loci associated with Sir2 or Hst1 using chromatin immunoprecipitation (ChIP). For *S. cerevisiae*, ChIP DNA was sequenced using high-throughput Illumina technology (ChIP-Seq). For *K. lactis*, ChIP DNA was hybridized to a tiled microarray (ChIP-chip). Next, we identified the genes whose transcription is influenced by Sir2 or Hst1 using Illumina sequencing of RNA (RNA-Seq). For this experiment, the cryptic mating-type loci were deleted so that loss of Sir2 would not lead to simultaneous expression of **a** and alpha transcription factors, a condition that specifies the diploid state and causes significant changes in gene expression. In *S. cerevisiae*, both *SIR2* and *HST1* were deleted in the same strain because these paralogs are known to substitute for one another (XIE *et al*. 1999; HICKMAN AND RUSCHE 2007). Genes directly regulated by Sir2 or Hst1 were defined as those that were associated with the deacetylase in the ChIP assay and increased at least twofold in the deletion strain compared to wild type.

In *S. cerevisiae*, 171 genes increased significantly in the *sir2Δ hst1Δ* strain. ScHst1 associated with 974 genes, of which 115 were significantly up-regulated in the *sir2Δ hst1Δ* strain (Figure 1A). Therefore, these 115 genes were identified as being directly regulated by ScHst1. Similarly, ScSir2 associated with a total of 176 genes, of which 10 also increased in expression and were thus identified as ScSir2-regulated (Figure 1B). In *K. lactis*, 1,159 genes were associated with KlSir2, and 255 genes increased in expression in the *sir2Δ* strain. 175 of these genes had both properties and were thus determined to be directly regulated by KlSir2 (Figure 1C). Most (153) of these 175 KlSir2-regulated genes were also upregulated in a separate *K. lactis* RNA-Seq dataset that was collected several years earlier (Figure S1). Unless otherwise noted, we focused on the 175 KlSir2-regulated genes (Figure 1C) that were identified based on the RNA-Seq conducted at the same time as the *S. cerevisiae* RNA-Seq. The ChIP and expression data for all annotated genes in both species are provided in Tables S3 and S4.

**Figure 1.**
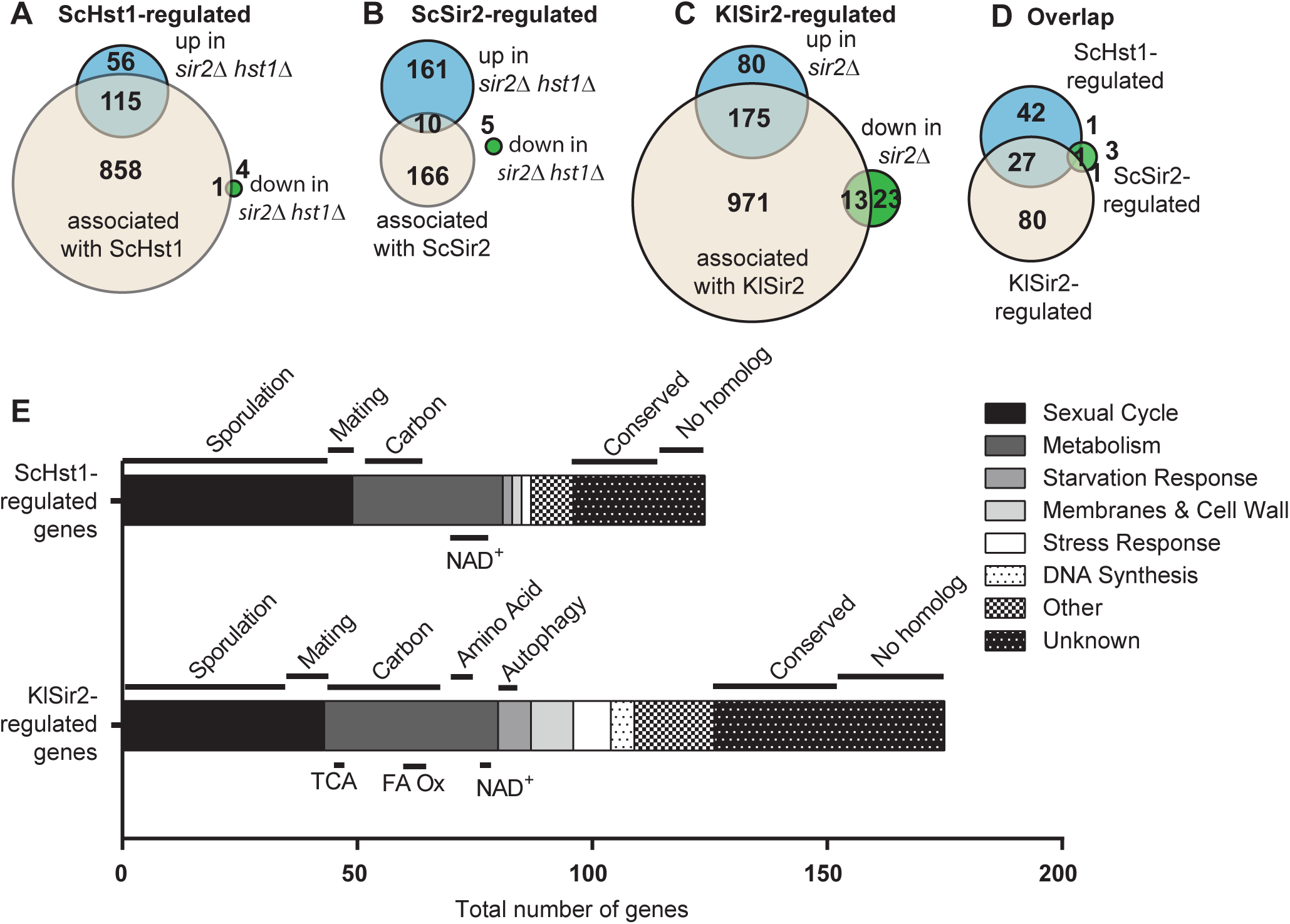
Identification of genes regulated by ScSir2, ScHst1, and KlSir2. (A) The overlap was determined for genes associated with ScHst1 (974), upregulated in *sir2Δ hst1Δ* cells (171), and downregulated in *sir2Δ hst1Δ* cells (5, including *SIR2* and *HST1*). (B) The overlap was determined for genes associated with ScSir2 (176), upregulated in *sir2Δ hst1Δ* cells (171), and downregulated in *sir2Δ hst1Δ* cells (5). (C) The overlap was determined for genes associated with KlSir2 (1159), upregulated in *sir2Δ* cells (255), and downregulated in *sir2Δ* cells (36, incluing *SIR2*). (D) The overlap was determined for genes regulated by ScSir2, ScHst1, and KlSir2. Of the 71 ScHst1-regulated genes with orthologs in *K. lactis*, 28 were also regulated by KlSir2. Of the six ScSir2-regulated genes with orthologs in *K. lactis*, two were also regulated by KlSir2. Of the 109 KlSir2-regulated genes with orthologs in *S. cerevisiae*, 28 were also regulated by ScHst1 and two were also regulated by ScSir2. (E) Genes regulated by ScHst1 and KlSir2 were manually grouped into functional categories based on GO terms and functional information (Tables S6 and S7). The bar graph represents the number of genes in each category.

ScSir2, ScHst1, and KlSir2 are all expected to repress transcription based on their deacetylase activity and previously described functions (RINE AND HERSKOWITZ 1987; XIE *et al*. 1999; HICKMAN AND RUSCHE 2009). Consistent with this expectation, genes associated with these deacetylases were more likely to be up-regulated than down-regulated in the absence of the deacetylase (Figure 1A-C). Indeed, in *S. cerevisiae* only one down-regulated gene was associated with ScHst1, and none were associated with ScSir2. However, in *K. lactis* 13 down-regulated genes were associated with KlSir2 (Figure 1C). To test statistically whether these genes are simply near KlSir2 peaks by chance or could be transcriptionally activated by KlSir2, we performed a Fisher’s exact test. We found a strong correlation between up-regulated genes and KlSir2 peaks (p < 2.2×10^-16^). In contrast, the correlation was less significant for down-regulated genes (p = 0.049). Therefore, KlSir2 functions primarily as a transcriptional repressor.

### Genes regulated by ScSir2 were adjacent to cryptic mating-type loci or telomeres

ScSir2 is thought to act primarily, if not exclusively, at the cryptic mating-type loci and telomeres (MARCHFELDER *et al*. 2003; ELLAHI *et al*. 2015). Consistent with this expectation, all but one of the ten genes we identified as ScSir2-regulated was near these loci (Table S5). Six genes were within 10 Kb of a telomere, and three genes were adjacent to the cryptic mating-type locus *HML*. No genes near *HMR* could be identified as ScSir2-repressed because *HMR* was deleted in the strains used for RNA-Seq. However, ScSir2 was associated with *HMR* in our ChIP-Seq study. The one ScSir2-regulated gene that was not near telomeres or mating-type loci was *YAT1*. This gene was also associated with ScHst1, and thus it is not clear which of these deacetylases is the primary regulator. We note that seven of the ten ScSir2-regulated genes were previously identified as being repressed by the Sir proteins (ELLAHI *et al*. 2015). We surveyed the functions of the ten ScSir2-regulated genes, but observed no common functional categories. We also compared the genome-wide distribution of ScSir2 observed in this study with data collected by two other labs. In agreement with (THURTLE AND RINE 2014), we found that ScSir2 is focused at telomeres and mating-type loci. In contrast, Sir2 was not associated with highly expressed genes (Figure S2), as reported (LI *et al*. 2013). We note that the ChIP-seq data that led to this conclusion had a low signal-to-noise ratio and many of the genes identified may represent a known “hyper-ChIP” artifact (TEYTELMAN *et al*. 2013). Therefore, our data support the previous understanding that ScSir2 performs a structural role at chromosome ends and maintains repression at the cryptic mating-type loci but does not regulate the expression of many genes. Consequently, perturbation of ScSir2 activity in low NAD^+^ is not expected to cause dramatic changes in gene expression.

### Genes regulated by ScHst1 function in the sexual cycle, NAD^+^ homeostasis, and scavenging for nutrients

In contrast to ScSir2-regulated genes, ScHst1-regulated genes were distributed throughout the genome. ScHst1 is known to repress genes involved in NAD^+^ biosynthesis and sporulation (XIE *et al*. 1999; BEDALOV *et al*. 2003), and indeed, these genes were well-represented among our 115 ScHst1-regulated genes (Figure 1E; Table S6). In particular, our list included five genes required for *de novo* synthesis of NAD^+^ and two that encode transporters of NAD^+^ precursors. It also included 43 genes involved in sporulation, most of which contribute to formation of the pro-spore membrane or spore wall. Moreover, 76 of the 115 genes (66%) are induced during sporulation (FRIEDLANDER *et al*. 2006; BORDE *et al*. 2009). Our data are consistent with previous studies, as 84 of our ScHst1-regulated genes (73%) were previously reported to be regulated by Hst1 or its DNA-binding partner Sum1 (BEDALOV *et al*. 2003; MCCORD *et al*. 2003).

We also observed additional groups of functionally-related genes that are regulated by ScHst1. For example, six genes are involved in cell fusion during mating and are associated with the shmoo tip, four genes encode hexose transporters, two genes are involved in allantoin degradation, two genes are involved in thiamine biosynthesis (LI *et al*. 2010), and two genes are involved in pyridoxine biosynthesis. Thus, when low NAD^+^ triggers the induction of ScHst1-regulated genes, *S. cerevisiae* responds by increasing the synthesis of NAD^+^ and other co-factors, scavenging for nutrients, and inducing genes required for mating and sporulation.

### Genes regulated by KlSir2 have additional functions not observed in S. cerevisiae

The functions of most KlSir2-regulated genes had to be inferred from their *S. cerevisiae* orthologs because few of these genes have been studied experimentally in *K. lactis*. We found that KlSir2-regulated genes have similar functions to ScHst1-regulated genes. In particular, two genes are involved in NAD^+^ homeostasis, 36 genes are involved in sporulation, nine genes are involved in mating, two genes are involved in allantoin metabolism, one gene is involved in thiamine biosynthesis, and one gene is involved in pyridoxine biosynthesis (Table S7). Thus, Sir2 has continued to regulate these biological processes over evolutionary time. However, we also found that a number of KlSir2-regulated genes were functionally distinct from ScHst1-regulated genes (Figure 1E). For example, KlSir2 regulates ten genes involved in metabolizing non-glucose carbon sources, four genes involved in DNA synthesis, and seven stress-response genes that mitigate the effects of heavy metals, oxidative stress, and DNA damage. Thus, when low NAD^+^ triggers the induction of KlSir2-regulated genes, *K. lactis* not only responds by increasing the level of NAD^+^ and by facilitating mating and sporulation, but it also mounts responses that buffer against stresses.

In *K. lactis*, the non-duplicated KlSir2 displays properties similar to both of its *S. cerevisiae* paralogs ScSir2 and ScHst1 (HICKMAN AND RUSCHE 2009). Therefore, we determined whether KlSir2-regulated genes are repressed in a manner similar to ScSir2, which acts with Sir4, or ScHst1, which acts with Sum1. Of the 175 KlSir2-regulated genes, 148 (84%) are repressed through a ScHst1-like mechanism, based on their increase in expression in the absence of KlSum1 (Table S7). However, only four are repressed by a ScSir2-like mechanism based on their increase in expression in the absence of KlSir4. Moreover, expression of KlSir2-regulated genes was correlated in *sir2Δ* and *sum1Δ* cells but not in *sir2Δ* and *sir4Δ* cells (Figure S3). Therefore, most KlSir2-regulated genes are regulated by the SUM1 complex, indicating that it is appropriate to compare these genes with those regulated by ScHst1.

### There was not much overlap between the gene sets regulated by ScHst1 and KlSir2

Given the functional overlap between the gene sets regulated by ScHst1 and KlSir2, it might be expected that the same genes in these functional categories would be regulated by these deacetylases in both species. However, only 29 genes were regulated in both species (Figure 1D), representing 16.5% of KlSir2-regulated genes and 24% of ScHst1-regulated genes. These genes included 17 genes involved in sporulation, one gene involved in mating, and one gene involved in NAD^+^ homeostasis. Thus, many of these common genes do participate in the biological processes that are regulated in both species. Nevertheless, many other genes associated with these processes are only regulated by Sir2 or Hst1 in one of the two species. This finding is consistent with a model in which the targets of the SUM1 complex shift over evolutionary time.

### ScHst1 and KlSir2 regulate NAD^+^ homeostasis through different genes

The regulation of NAD^+^ biosynthesis by ScHst1 has been described as a feedback loop (BEDALOV *et al*. 2003). In particular, a drop in intracellular NAD^+^ levels would reduce the activity of the NAD^+^-dependent deacetylase ScHst1, relieving its repression of genes that boost NAD^+^ levels. These genes include those involved in *de novo* NAD^+^ biosynthesis (*BNA* genes) and the import of NAD^+^ precursors (*TNA1* and *NRT1*). We also observed that these genes are regulated by ScHst1, and we found that a similar feedback loop exists for KlSir2 (Figure 2, Tables S6 and S7). However, only one gene, the transporter *TNA1*, was regulated in both species. *K. lactis* lacks the genes for *de novo* biosynthesis, precluding them from regulation by KlSir2. In addition, *K. lactis* lacks a unique transporter for the NAD^+^ precursor nicotinamide riboside. A single *K. lactis* gene, KLLA0D00550g, is related to three paralogous *S. cerevisiae* transporters, including the nicotinamide riboside transporter *NRT1* and two thiamine transporters. This arrangement may account for KlSir2 instead regulating enzymes that process NAD^+^ precursors. In summary, both species use an NAD^+^-dependent deacetylase as part of a feedback mechanism to maintain NAD^+^ levels, but the particular genes involved in this circuit are different. This finding suggests that even though the specific targets of Sir2 have shifted over evolutionary time, it has maintained regulation of NAD^+^ homeostasis. This finding also supports the notion that Sir2 and Hst1 are sensors that tune gene expression in response to fluctuations in intracellular NAD^+^ levels.

**Figure 2.**
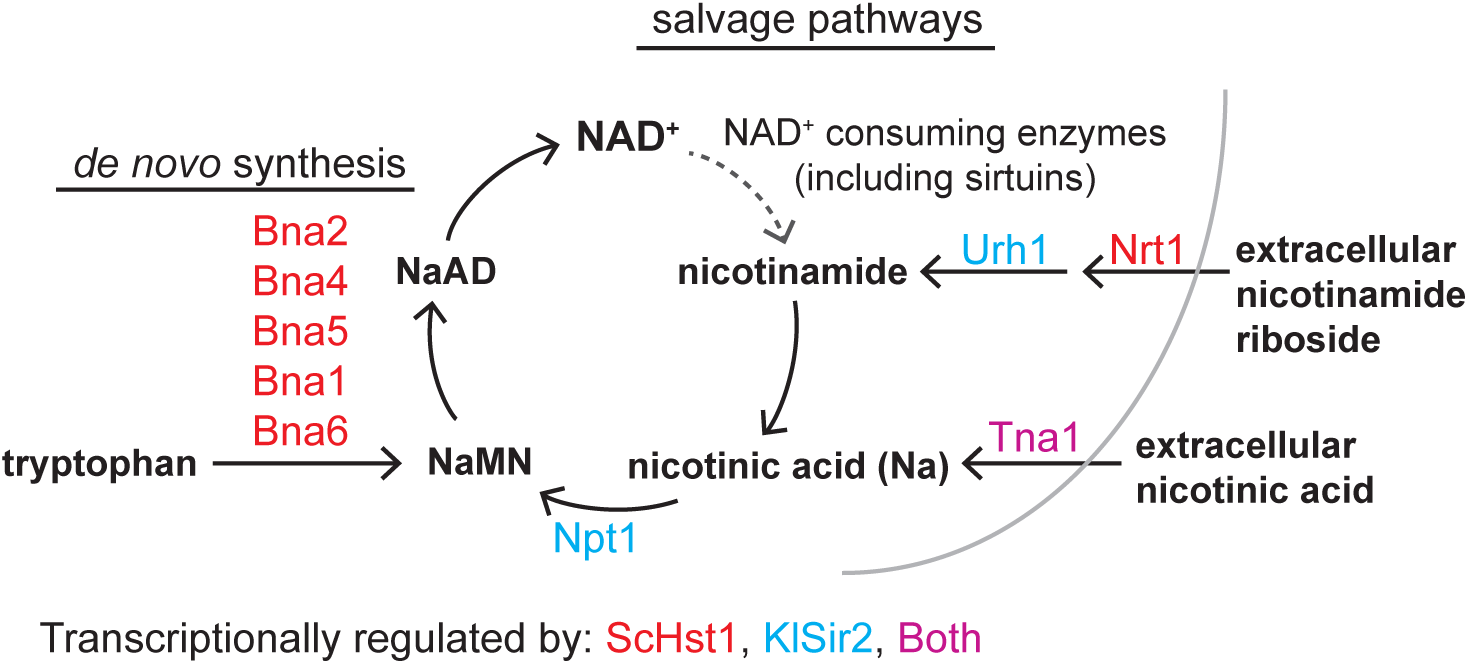
ScHst1 and KlSir2 regulate NAD^+^ homeostasis through different genes. The NAD^+^ *de novo* biosynthesis and salvage pathways are shown. Genes regulated by ScHst1 are colored red, by KlSir2 are colored blue, and by both ScHst1 and KlSir2 are purple. *URH1* was only upregulated in the older RNA-Seq dataset. NAD^+^ is nicotinamide adenine dinucleotide, NaMN is nicotinic acid mononucleotide, NaAD is nicotinic acid adenine dinucleotide.

### KlSir2 regulates genes involved in utilization of carbon sources other than glucose

We observed that genes involved in carbon metabolism were regulated by both ScHst1 and KlSir2 (Figure 3, Tables S6-S8). Many of these genes facilitate growth in the absence of glucose. For example, some genes metabolize sugars other than glucose, such as galactose or lactose. Other genes metabolize non-sugars such ethanol, glycerol, fatty acids, or amino acids. We also observed regulation of genes involved in the TCA cycle and the related glyoxylate and methyl citrate cycles. The TCA cycle is required for aerobic respiration and is an alternative to fermentation, the preferred way for *S. cerevisiae* to process glucose. The methylcitrate cycle is a variation of the TCA cycle in which the three-carbon compound propionyl-CoA is metabolized in place of the two-carbon compound acetyl-CoA. This cycle enables metabolism of propionate and fatty acids with odd numbers of carbons. The glyoxylate cycle is also a variation of the TCA cycle, in which the steps that produce CO_2_ are bypassed. Instead, the carbon atoms are shunted into gluconeogenesis, allowing a cell to build sugars from acetyl groups. Doing so is necessary for synthesizing nucleotides and cell wall carbohydrates when sugars are not available in the environment.

**Figure 3.**
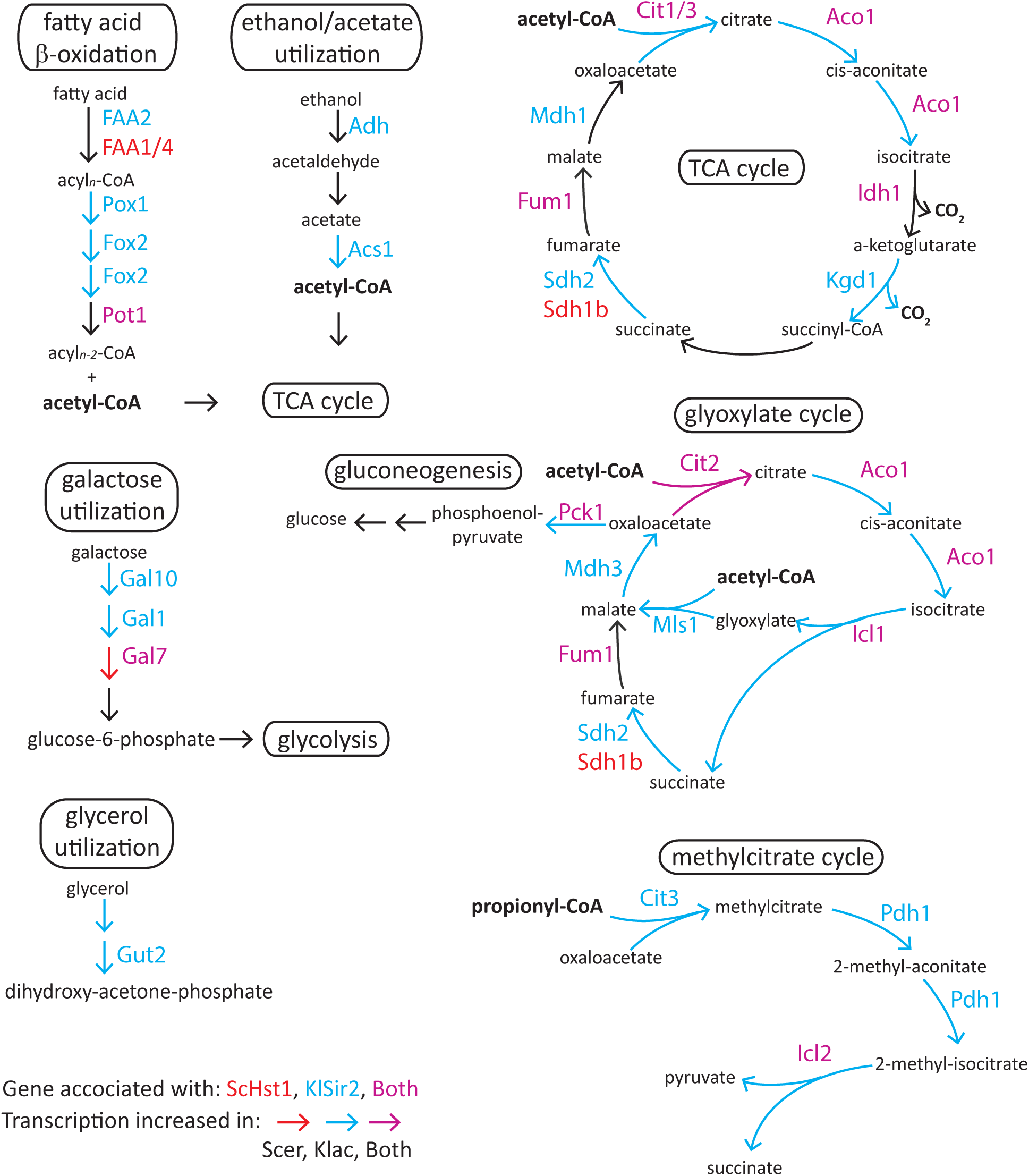
Metabolic pathways regulated by ScHst1 and KlSir2. Regulation by ScHst1 and KlSir2 was evaluated for each gene in metabolic pathways of interest. Gene names are colored red for association with ScHst1, blue for association with KlSir2, and purple for association with both sirtuins. Arrows are colored red if the gene increased in *hst1Δ sir2Δ* compared to wild-type *S. cerevisiae*, blue if the gene increased in *sir2Δ* compared to wild-type *K. lactis*, and purple if the gene increased in both species. For *K. lactis*, induction was evaluated using both RNA-Seq datasets. The genes evaluated are listed in Table S9. Galactose is converted to glucose-6-phosphate and then enters glycolysis. Glycerol is converted to dihydroxy-acetone-phosphate, an intermediate in glycolysis and gluconeogenesis. Non-fermentable carbon sources, including fatty acids, ethanol, and acetate, are metabolized to acetyl-CoA via the fatty acid β-oxidation and ethanol/acetate utilization pathways. The acetyl-CoA then feeds into the TCA cycle to produce energy or the glyoxylate cycle and gluconeogenesis to produce glucose. Fatty acids with odd numbers of carbons also generate propionyl-CoA, which is metabolized via the methylcitrate cycle.

To assess the extent to which metabolic pathways are regulated by ScHst1 and KlSir2, we scored all the genes in each pathway of interest for the association of ScHst1 or KlSir2 and for induction in the absence of the deacetylase (Figure 3, Table S9). We found some pathways are regulated by KlSir2 but not ScHst1, including the methylcitrate, fatty acid β-oxidation, glycerol utilization, and ethanol/acetate utilization pathways. In contrast, both sirtuins have the potential to regulate the TCA and glyoxylate cycles. Thus, in *K. lactis* a drop in intracellular NAD^+^ that compromises KlSir2 activity would lead to the increased expression of genes required to utilize non-sugar carbon sources.

Given that metabolic flux in *K. lactis sir2Δ* cells might be shifted away from glucose consumption, we compared the growth of wild-type and *sir2Δ* strains in several carbon sources. In glucose, *K. lactis sir2Δ* cells grew more slowly than wild-type cells (Figure 4A). In contrast, in *S. cerevisiae*, loss of Sir2 and Hst1 did not affect growth in glucose (Figure 4C). Interestingly, the *K*. *lactis sir2Δ* cells actually grew faster than wild-type cells in glycerol (Figure 4B). For *S. cerevisiae*, growth did not occur in minimal medium with 3% glycerol. These results are consistent with the model that KlSir2 promotes the ability of *K. lactis* cells to utilize glucose efficiently by damping down pathways that favor other carbon sources. Such repression is presumably relieved in low NAD^+^.

**Fig 4.**
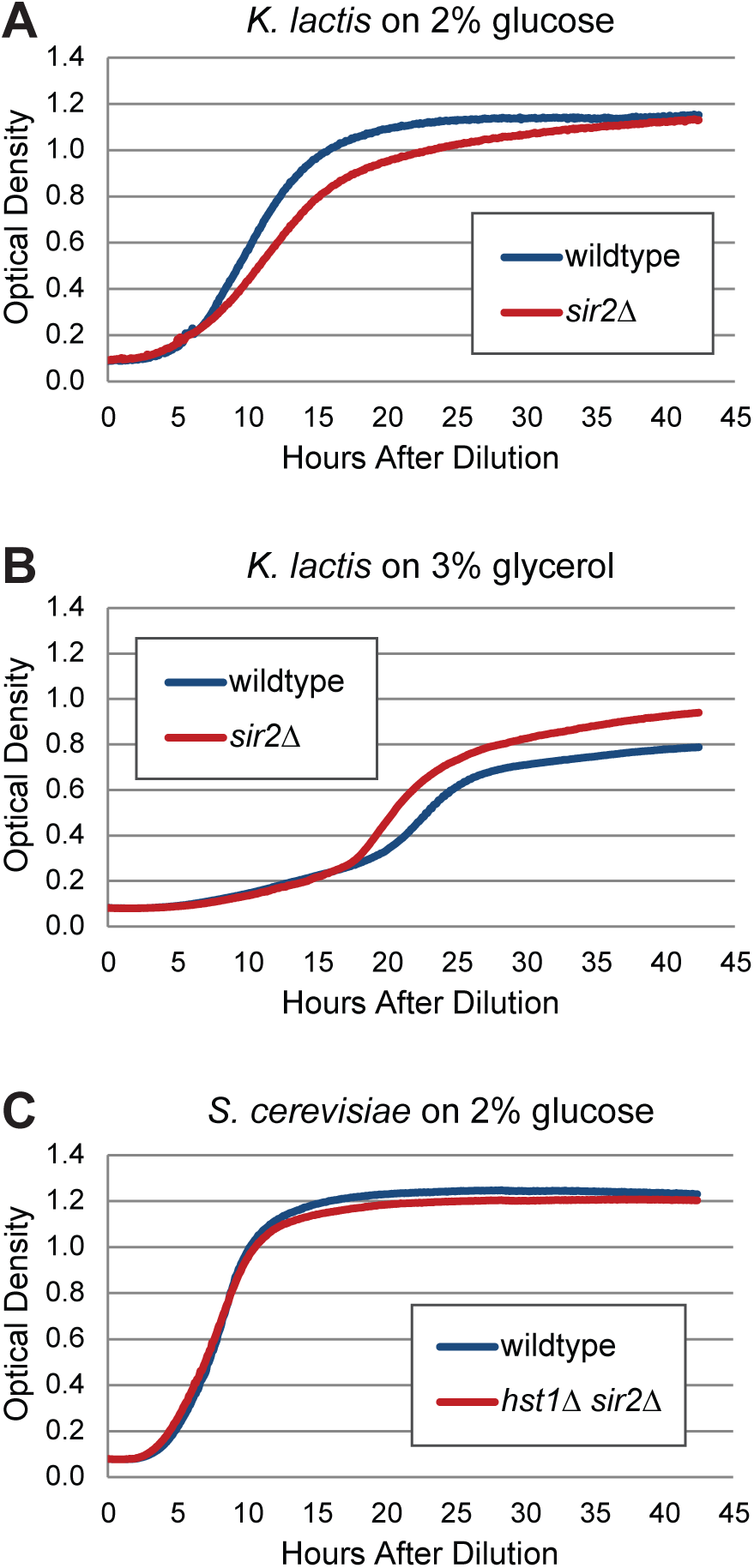
KlSir2 but not ScHst1 influenced growth rate in minimal medium. (A) *K. lactis* cells with or without SIR2 were grown in minimal medium (YM) with 2% glucose, and the density of the culture was recorded over time using a microplate reader. *K. lactis* strains were *SIR2* (LRY2992) and *sir2Δ* (LRY2993). (B) *K. lactis* cells were grown in minimal medium with 3% glycerol. (C) *S. cerevisiae* cells with or without *SIR2* and *HST1* were grown in minimal medium with 2% glucose. The *S. cerevisiae* cells did not grow in minimal medium with 3% glycerol. *S. cerevisiae* strains were *HST1 SIR2* (LRY3093) and *hst1Δ sir2Δ* (LRY3099).

### KlSir2 impacts resistance to heavy metals

Several genes regulated by KlSir2 are involved in stress responses (Table S7), including three genes responsible for removing the heavy metal arsenic from cells. In contrast, ScHst1 and ScSir2 do not regulate these or similar genes. To test whether the presence of Sir2 or Hst1 affects resistance to arsenic, we grew wild-type and mutant cells on YPD supplemented with sodium arsenate. We found that growth of wild-type *K. lactis* cells was reduced in sodium arsenate, whereas *sir2Δ* cells grew similarly in the presence or absence of sodium arsenate (Figure 5A). Note that in this medium containing glucose, the *sir2Δ* cells grew more slowly than wildtype cells, consistent with our findings in Figure 4. *S. cerevisiae* cells were more resistant to arsenate than *K. lactis*, as a higher concentration was required to impact growth. In contrast to *K. lactis*, wildtype and *sir2Δ hst1Δ S. cerevisiae* cells grew similarly in sodium arsenate (Figure 5B). Thus, KlSir2 impedes the ability of *K. lactis* cells to detoxify arsenic, but ScHst1 and ScSir2 do not act similarly in *S. cerevisiae*.

**Figure 5.**
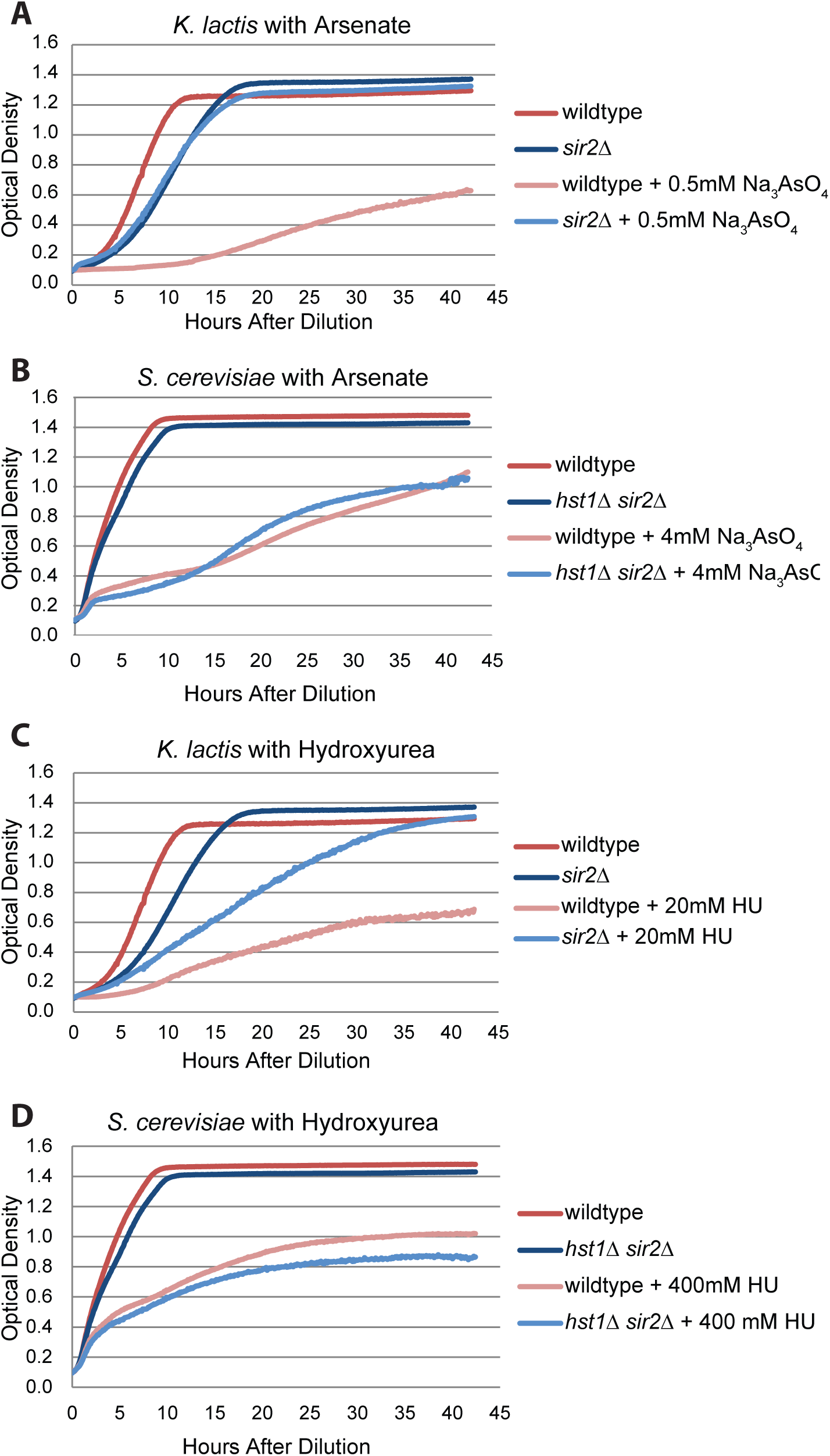
KlSir2 but not ScHst1 influenced growth on arsenic and hydroxyurea. (A) *K. lactis* cells with and without *SIR2* were grown in rich medium (YPD) alone or with 0.5 mM sodium arsenate, and the density of the culture was recorded over time using a microplate reader. The same strains were used as for Figure 4. (B) *S. cerevisiae* cells were grown in YPD alone or with 4 mM sodium arsenate, as described for panel A. (C) *K. lactis* cells were grown in YPD alone or with 20 mM hydroxyurea. (D) *S. cerevisiae* cells were grown in YPD alone or with 400 mM hydroxyurea.

### KlSir2 impacts resistance to hydroxyurea, which depletes dNTPs

Several genes regulated by KlSir2 are involved in DNA synthesis (Table S7), including the gene encoding ribonucleotide reductase, the enzyme that reduces ribonucleotides (NTPs) to deoxyribonucleotides (dNTPs). In contrast, ScHst1 and ScSir2 do not regulate these genes. We assessed whether Sir2 or Hst1 affects growth of cells in the presence of an inhibitor of ribonucleotide reductase, hydroxyurea (HU). We found that when *K. lactis* cells were grown in 20 mM HU, the growth was more severely impacted for wildtype cells than *sir2Δ* cells (Figure 5C). *S. cerevisiae* cells were more resistant to HU than *K. lactis*, as a higher concentration was required to impact growth. Nevertheless, wildtype and *sir2Δ hst1Δ S. cerevisiae* cells grew similarly in 400 mM HU (Figure 5D). Therefore, KlSir2, but not ScHst1 or ScSir2, influenced the ability of cells to grow when ribonucleotide reductase is partially inhibited. These results are consistent with KlSir2 reducing the production of dNTPs.

### Genes regulated by ScHst1 and KlSir2 are less likely than other genes to be evolutionarily conserved

The results above reveal that *K. lactis* and *S. cerevisiae* have distinct growth phenotypes that can be attributed to genes that are repressed by KlSir2 but not ScHst1. In addition to these conserved genes that are only regulated by the SUM1 complex in *K. lactis*, we found that a disproportionately high number of regulated genes only occur in one of the two species. To identify orthologs, we used a two-way BLASTP procedure described in the methods. This approach allowed us to assign *S. cerevisiae* orthologs for 103 of the 175 KlSir2-regulated genes. Of the remaining genes, 56 had no significant BLASTP hit in *S. cerevisiae*. Another 16 had a hit but the two-way BLASTP search did not return to the starting *K. lactis* gene, indicating that the *S. cerevisiae* hit is actually the homolog of a different *K. lactis* gene. Thus, we identified *S. cerevisiae* orthologs for just 103 (59%) of the genes regulated by KlSir2. (Manual refinement ultimately identified orthologs for 109 (62%) genes.) In contrast, for the genome as a whole, the same analysis identified *S. cerevisiae* orthologs for 90% of all *K. lactis* genes. To estimate the statistical probability that a random set of genes would deviate so much from the genome-wide percentage, we generated 10,000 random sets of 175 *K. lactis* genes and recorded the percentage of genes in each set that had *S. cerevisiae* orthologs. As expected, the results are distributed around the genome-wide value of 90% (Figure 6A). Importantly, none of the 10,000 trials resulted in a percentage close to 59%, indicating that random chance does not explain the low percentage of KlSir2-regulated genes with *S. cerevisiae* orthologs. We extended this analysis to other species and found that KlSir2-regulated genes are also less likely to have orthologs in nine other fungal species (Figure 6B). Therefore the set of genes regulated by KlSir2 is enriched for genes that are not widely conserved. We also found the same trend for ScHst1-regulated genes, with regulated genes having a lower percentage of orthologs in other species compared to total *S. cerevisiae* genes (Figure 6C). Thus, genes regulated by Sir2 and Hst1 are more likely to be species-restricted than the average fungal gene. This observation is consistent with the model that sirtuins regulate genes that result in species-appropriate responses to low NAD^+^.

**Figure 6.**
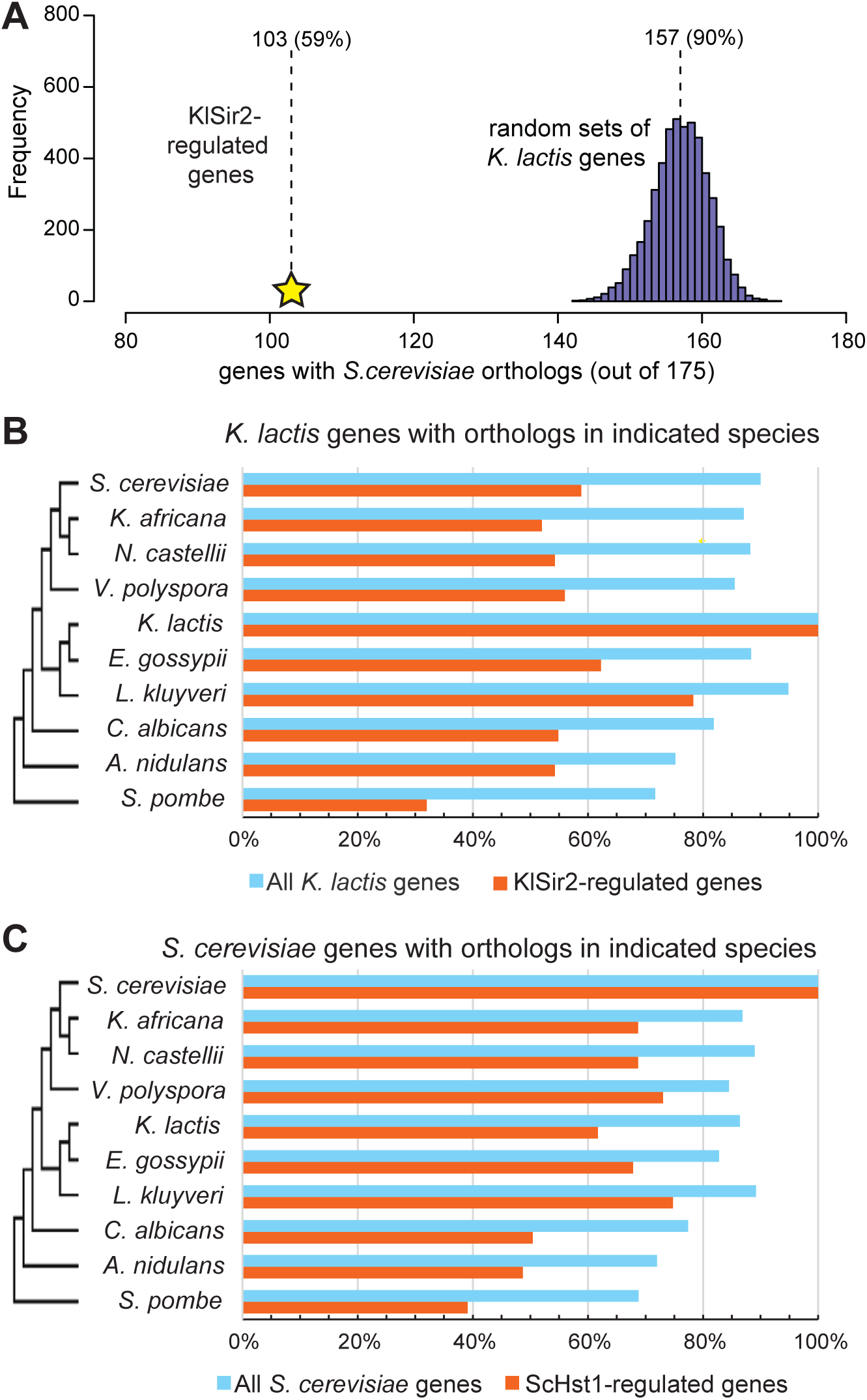
Genes regulated by ScHst1 and KlSir2 were less likely than other genes to be widely conserved. (A) 10,000 sets of 175 randomly selected *K. lactis* genes were evaluated for the number of genes with *S. cerevisiae* orthologs. The results were distributed around 157 genes (90%). For the 175 KlSir2-regulated genes, 103 (59%) had *S. cerevisiae* orthologs. This number is well outside the distribution for the randomly selected genes. (B) For *K. lactis*, the percentage of genes with orthologs in each of nine species is graphed for the 5076 total genes (blue) and the 175 KlSir2-regulated genes (red). (C) For *S. cerevisiae*, the percentage of genes with orthologs in each of nine species is graphed for the 5906 total genes (blue) and the 115 ScHst1-regulated genes (red).

### KlSir2 regulates synthesis of pulcherrimin, an iron-chelating compound not produced by most yeast

Two KlSir2-regulated genes that are not found in *S. cerevisiae* or many other yeasts are *PUL1* and *PUL2*. Together, Pul1 and Pul2 synthesize the secreted siderophore pulcherriminic acid, which chelates iron(III) to form a red-colored compound pulcherrimin (KRAUSE *et al*. 2018). *K. lactis* scavenges iron by importing pulcherrimin via a specific transporter, Pul3. It is speculated that microbes that use pulcherrimin to sequester iron gain an advantage over microbes that lack the transporter (SIPICZKI 2006; ORO *et al*. 2014; KRAUSE *et al*. 2018). To determine whether KlSir2 influences the production of pulcherrimin, we spotted wild-type and *sir2Δ* cells on rich medium (YPD) supplemented with FeCl_3_ (Figure 7). After two days, we observed a red halo surrounding the *K. lactis* cells. This halo was likely due to pulcherrimin, as it did not form on plates lacking FeCl_3._ Moreover, it did not form around *S. cerevisiae* cells, which lack *PUL1* and *PUL2*. Importantly, in *sir2Δ K. lactis* cells, the red halo was more intense in color, consistent with KlSir2 suppressing the production of pulcherrimin.

**Figure 7.**
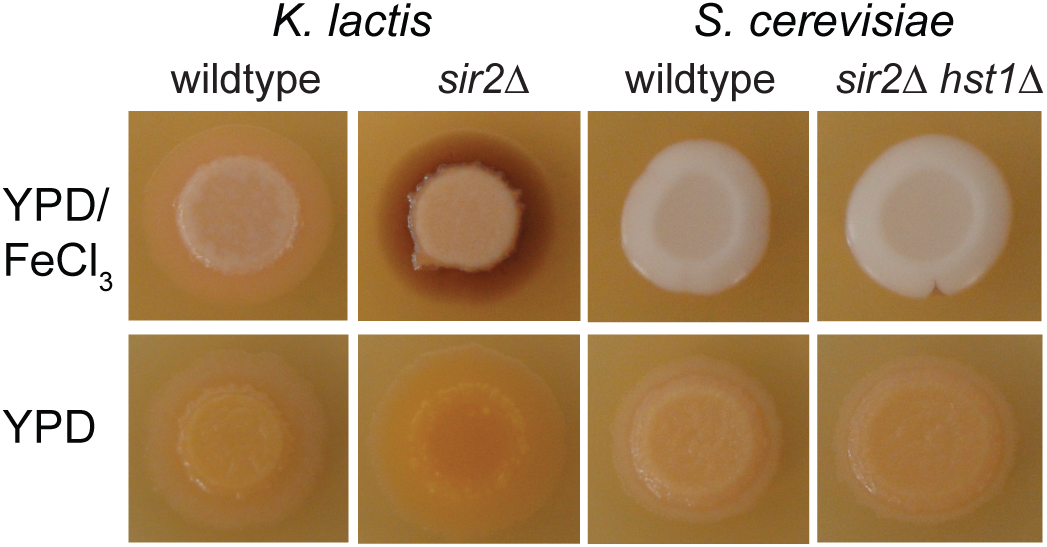
KlSir2 influenced production of the siderophore pulcherrimin. Exponentially growing cells were collected and spotted on rich medium (YPD) with or without 3.7 mM FeCl_3_. The plates were incubated two days and then imaged. The same strains were used as for Figure 4.

### Loss of Hst1 and Sir2 reduced mating in both S. cerevisiae and K. lactis

We did observe some functional categories of genes that are regulated by both ScHst1 and KlSir2, including genes required for mating. In *S. cerevisiae*, these genes act relatively late in the mating pathway (Table S6). Specifically, four of the six mating genes encode proteins that are localized to the shmoo tip and are involved in cell fusion. Another gene is involved in nuclear fusion, and the last damps down pheromone signaling. Similarly, three KlSir2-regulated genes also encode proteins associated with the shmoo tip and cell fusion. However, there are also four KlSir2-regulated genes involved in the earliest steps of (Table S7). These include a pheromone (alpha factor) and two subunits of the G protein that signals pheromone binding. Given that *K. lactis* delays mating until it encounters nutrient deprivation, an appealing hypothesis is that nutrient deprivation is associated with a drop in intracellular NAD^+^, which in turn triggers a reduction of KlSir2 activity and increased expression of mating genes. Thus, KlSir2 could help restrict mating to nutrient poor conditions. In contrast, ScHst1 may not play this role, as *S. cerevisiae* mates in rich nutrient conditions.

To test the hypothesis that KlSir2 hinders mating by repressing mating genes, we developed a quantitative mating assay based on imaging cytometry. This approach was necessary because *K. lactis* mates at a low frequency. For both wild type and *sir2Δ* strains, we generated *MAT**a*** cells that expressed GFP and *MATα* cells that expressed mCherry. In addition, the cryptic mating-type loci were deleted so *sir2Δ* cells would not be sterile pseudodiploids, and the strains were made prototrophic to eliminate nutritional dependencies that might influence mating. *MAT**a*** and *MATα* haploid cells were grown together on malt extract to induce mating and were then examined using an Image Stream cytometer. Mated zygotes were identified as those cells that were both red and green and had the classic hourglass shape. Consistent with previous studies (HERMAN AND ROMAN 1966), the efficiency of mating was very low. Nevertheless, zygote formation was significantly lower in the *sir2Δ* strain, compared to the wild type (Figure 8A). This result indicated that although Sir2 does influence the efficiency of mating, the direction of change was opposite to our prediction that increased expression of mating genes in the *sir2Δ* strain would enhance mating.

**Figure 8.**
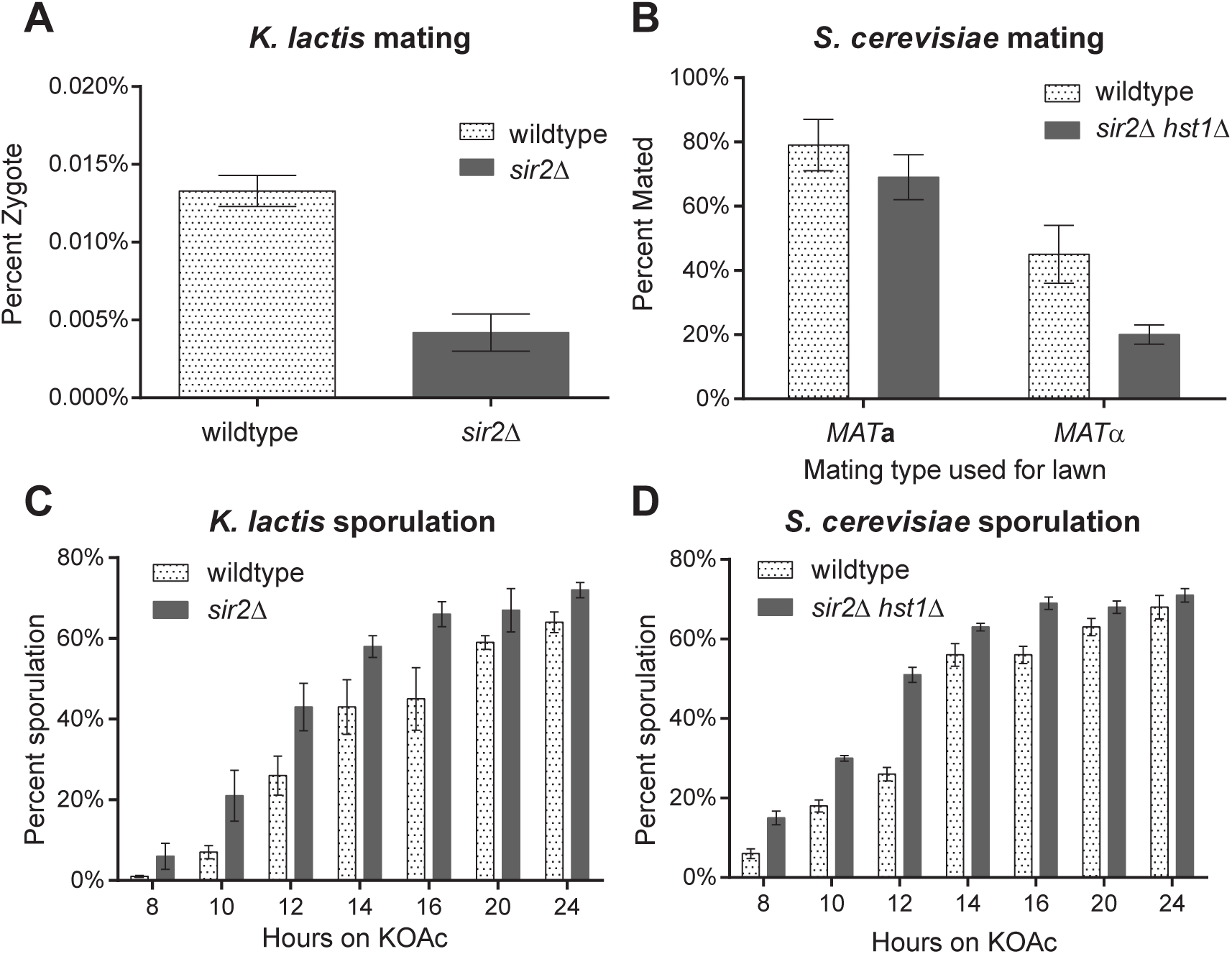
Loss of ScHst1 and KlSir2 decreased mating and increased sporulation. (A) Percentage of observed *K. lactis* cells that were zygotes. *MAT**a*** and *MATα* cells were co-cultured on malt extract for three days, resuspended in water, sonicated, and then examined using ImageStream cytometry. For each sample,100,000 events were collected. Cells that were in focus and showed both red and green fluorescence were identified using IDEAS software (EMD Millipore). These cells were then examined manually for an hourglass appearance typical of zygotes. Strains were *MAT**a** SIR2* mCherry (LRY3037 and 3038), *MAT**a** sir2Δ* mCherry (LRY3039 and 3040), *MATα SIR2* GFP (LRY3053 and 3054), and *MATα sir2Δ* GFP (LRY3055 and 3056). All strains lacked *HML* and *HMR*. (B) Percentage of *S. cerevisiae* haploids that mated to produce diploids. A lawn of one mating-type was prepared using 2 OD of cells spread on minimal plates supplemented with histidine, tryptophan, and leucine. The other mating-type (2×10^-5^ OD) was spread over the lawn to determine the number of cells that could mate. Only diploid cells could grow on these selective plates. The same volume of cells was also spread on rich (YPD) plates to determine the number of viable cells. Strains were *MAT***a** *HST1 SIR2 hmlαΔ* (LRY3104), *MAT***a** *hst1Δ sir2Δ hmlαΔ* (LRY3101), *MATα HST1 SIR2 hmraΔ* (LRY3093), and *MATα hst1Δ sir2Δ hmraΔ* (LRY3099). (C) Percent sporulation over time for wild-type and *sir2Δ K. lactis* cells. Freshly grown diploid cells were patched onto KOAc sporulation plates and incubated at 30°C. At two hour intervals, cells were resuspended in water and examined microscopically. All cells in three fields of vision were scored as either having sporulated (tetrad morphology) or not. The experiment was conducted twice with four biological replicates (diploid strains) each time. Strains used were *SIR2* (LRY3019, 3029-31) and *sir2Δ* (LRY3020, 3032-34). (D) Percent sporulation for wild-type and *hst1Δ sir2Δ S. cerevisiae* cells. The assay was conducted as described for panel B. Strains used were *HST1 SIR2* (LRY3108-3111) and *hst1Δ sir2Δ* (LRY3112-3115).

For comparison, we also examined whether the absence of ScHst1 and ScSir2 influenced the ability of *S. cerevisiae* cells to mate. In this case, it was not necessary to use imaging cytometry because the efficiency of mating in *S. cerevisiae* is much higher. Instead, a known number of haploid cells of one mating-type was spread on a lawn of the opposite mating type. Only diploids had the necessary markers to grow on the selective plate, allowing us to determine the fraction of cells that mated. As for *K. lactis*, the silent mating-type loci were deleted so the loss of Sir2-silencing would not lead to a sterile pseudodiploid state. The *sir2Δ hst1Δ* strains had slightly reduced mating when the *MAT**a*** stain was used as the lawn and a more pronounced decrease in mating when the *MATα* stain was used as the lawn (Figure 8B). Thus, the loss of ScSir2 and ScHst1 reduced mating in *S. cerevisiae*, just as the loss of KlSir2 did in *K. lactis*. Therefore, even though only KlSir2 regulates early mating genes, ScHst1 and KlSir2 both impact the mating process similarly. We speculate that the timing of mating events may be perturbed in the absence of ScHst1 or KlSir2.

### Loss of Hst1 and Sir2 shortened the time to sporulation in both S. cerevisiae and K. lactis

When diploid yeast cells are starved for nitrogen, they initiate meiosis. The four resulting haploid nuclei are each encased in a special cell wall and become spores. Both ScHst1 and KlSir2 repress sporulation genes (Figure 1E) (XIE *et al*. 1999; HICKMAN AND RUSCHE 2009). However, different classes of genes are regulated in the two species. In *S. cerevisiae*, 84% (36/43) of ScHst1-regulated sporulation genes are “mid-sporulation genes” involved in formation of the spore membrane and wall. In contrast, in *K. lactis* only 64% (23/36) of KlSir2-regulated genes are involved in this phase of sporulation. Other genes are involved in earlier steps of meiosis, including four genes involved in chromosome pairing and segregation. It is particularly striking that KlSir2 regulates three meiotic transcription factors, including *IME1*, the master inducer of meiosis (Table S7). Based on this observation, we hypothesized that loss of KlSir2 would advance the timing of sporulation in *K. lactis* whereas loss of ScHst1 would not do so in *S. cerevisiae*. To test this hypothesis, diploid cells freshly grown from freezer stocks were placed on sporulation medium. These cells were examined microscopically every two hours, and the fraction of tetrads (products of sporulation) was scored. As predicted, we found that at each time point a greater percentage of *K. lactis sir2Δ* cells had sporulated compared to wild-type cells (Figure 8C). Surprisingly however, we found the same trend for *S. cerevisiae* cells (Figure 8D). Therefore, both KlSir2 and ScHst1 delay sporulation. Even though KlSir2 regulates more early-sporulation genes than ScHst1 does, both deacetylases impact the sporulation process similarly.

## DISCUSSION

In this study, we examined the hypothesis that Sir2 functions as a transcriptional rewiring point, potentially leading to species-appropriate adaptive responses to conditions that decrease intracellular NAD^+^ levels. We compared genes regulated by Sir2 and its paralog Hst1 in two yeast species that diverged over 100 million years ago (SHEN *et al*. 2018), and we found that some biological processes regulated by these deacetylases are common to both species. Nevertheless, the specific genes that are regulated are distinct, indicating significant plasticity in the targets of Sir2 and Hst1 over evolutionary time. In addition, KlSir2 regulates genes in functional categories not regulated by ScHst1 or ScSir2, such as utilization of non-glucose carbon sources, DNA replication, resistance to heavy metals, and production of the siderophore pulcherrimin. These findings indicate the Sir2 can serve as a transcriptional rewiring point.

It is striking that even though *S. cerevisiae* and *K. lactis* have evolved separately for over 100 million years, many of the same biological processes are regulated by Sir2 and Hst1 in the two species. These processes include NAD^+^ homeostasis, mating, and sporulation. This finding suggests that the last common ancestor of *S. cerevisiae* and *K. lactis* also employed Sir2 to regulate these processes and that connecting these processes to NAD^+^ levels has remained evolutionarily advantageous. Indeed, there is a clear benefit to regulating NAD^+^ homeostasis genes through a feedback loop in which a drop in NAD^+^ levels relieves repression of these genes. It may also be advantageous for mating and sporulation to be regulated by an NAD^+^-dependent repressor, as these events often occur under low nutrient conditions that could coincide with decreased availability of NAD^+^.

It is also striking that different genes involved in the same biological processes are regulated by ScHst1 and KlSir2. For example, only 17 out of 43 ScHst1-regulated sporulation genes (40%) are also regulated in *K. lactis*. This finding could indicate that it is not critical which specific genes within a functional category are regulated. Alternatively, it could indicate nuanced differences in how the two species mount developmental programs such as mating or sporulation. For example, it might be more advantageous for *K. lactis* than *S. cerevisiae* to employ Sir2 to integrate information about NAD^+^ levels into the expression of early sporulation genes, including the master regulator *IME1*. If so, there might be particular situations in which fluctuations in NAD^+^ levels would impact mating or sporulation in one species but not the other. Nevertheless, under the conditions we examined, the loss of the Sir2 and Hst1 impacted mating and sporulation similarly in both species.

An important finding is that KlSir2 regulates additional genes not associated with the functional categories regulated by ScHst1. Induction of these genes in low NAD^+^ conditions might be adaptive for *K. lactis* but not for *S. cerevisiae*. For example, KlSir2 regulates genes that metabolize non-glucose carbon sources. The regulation of these genes in *K. lactis* might relate to its use of respiration rather than fermentation in the presence of oxygen. We also observed that KlSir2 regulates genes involved in heavy metal efflux, dNTP production, and siderophore synthesis, whereas these genes were not regulated in *S. cerevisiae*. Similarly, others have found that in the pathogenic yeast *Candida glabrata*, Sir2 and Hst1 regulate genes that favor growth in a mammalian host (DE LAS PENAS *et al*. 2003; ORTA-ZAVALZA *et al*. 2013). These observations suggest that Sir2 does serve as a rewiring point, such that some processes are linked to NAD^+^ availability only in certain species.

It is notable that a higher than expected number of Sir2- and Hst1-regulated genes lack orthologs in other fungal species. This observation is consistent with sirtuin deacetylases contributing to the acquisition of distinct responses to low NAD^+^. Only 59% of KlSir2-regulated genes have *S. cerevisiae* orthologs, whereas 90% of all *K. lactis* genes do. Similarly, 62% of ScHst1-regulated genes compared to 86% of all *S. cerevisiae* genes have *K. lactis* orthologs. Thus, Sir2- and Hst1-repressed genes are more likely than the average gene to be restricted to a few species. Such genes are likely to provide unique functions, such as siderophore production, to the species in which they reside, and hence their being regulated by Sir2 or Hst1 is consistent with the hypothesis that bringing new processes under the control of sirtuins is associated with organism-specific responses to low NAD^+^ stress. In the future, it will be interesting to determine the functions of these species-restricted genes.

An important technical consideration that emerged during this study is that some regulated genes were likely missed by the approach of combining ChIP and gene expression data. In particular, some Sir2- and Hst1-repressed genes might require a transcriptional activator for expression but that activator might not have been available under the standard growth conditions we used. Consequently, such a gene would not be induced in the absence of the sirtuin repressor. This scenario could account for the larger number of genes that were associated with Sir2 or Hst1 than were induced in the absence of these sirtuins. It is therefore probable that under other growth conditions, additional Sir2- and Hst1-regulated genes could be identified.

In summary, our results are consistent with the hypothesis that sirtuins are rewiring points that allow species to evolve distinct responses to low NAD^+^ stress. Because sirtuins require NAD^+^ for enzymatic activity, they are hard-wired to respond to fluctuations in intracellular NAD^+^ and can be thought of as dedicated rewiring points for making a cellular process sensitive to NAD^+^ levels. Bringing new genes under the control of Sir2 or Hst1 enables yeast species to develop patterns of gene expression that evoke an appropriate response to low NAD^+^, potentially increasing fitness.

## ACKNOWLEDGEMENTS

We thank David MacAlpine for assistance designing the *K. lactis* microarrays. We thank Stefan Astrom, Juergen Heinisch, Peter Philippsen, Lorraine Pillus, Jasper Rine, Hana Sychrova, and Christopher Taron (New England Biolabs) for strains and plasmids. We thank the members of the Rusche lab for suggestions and support.

## Author Contributions

The *K. lactis* ChIP-chip dataset was generated by MH. The *K. lactis* 2012 RNA-seq dataset was generated by SH. The experiments shown in Figures 4, 5, and 7 were conducted by HM. All other experiments were conducted by KH. The bioinformatics analysis was performed by LZ and TL. The manuscript and figures were prepared by KH and LR.

---

**Figure S1.**
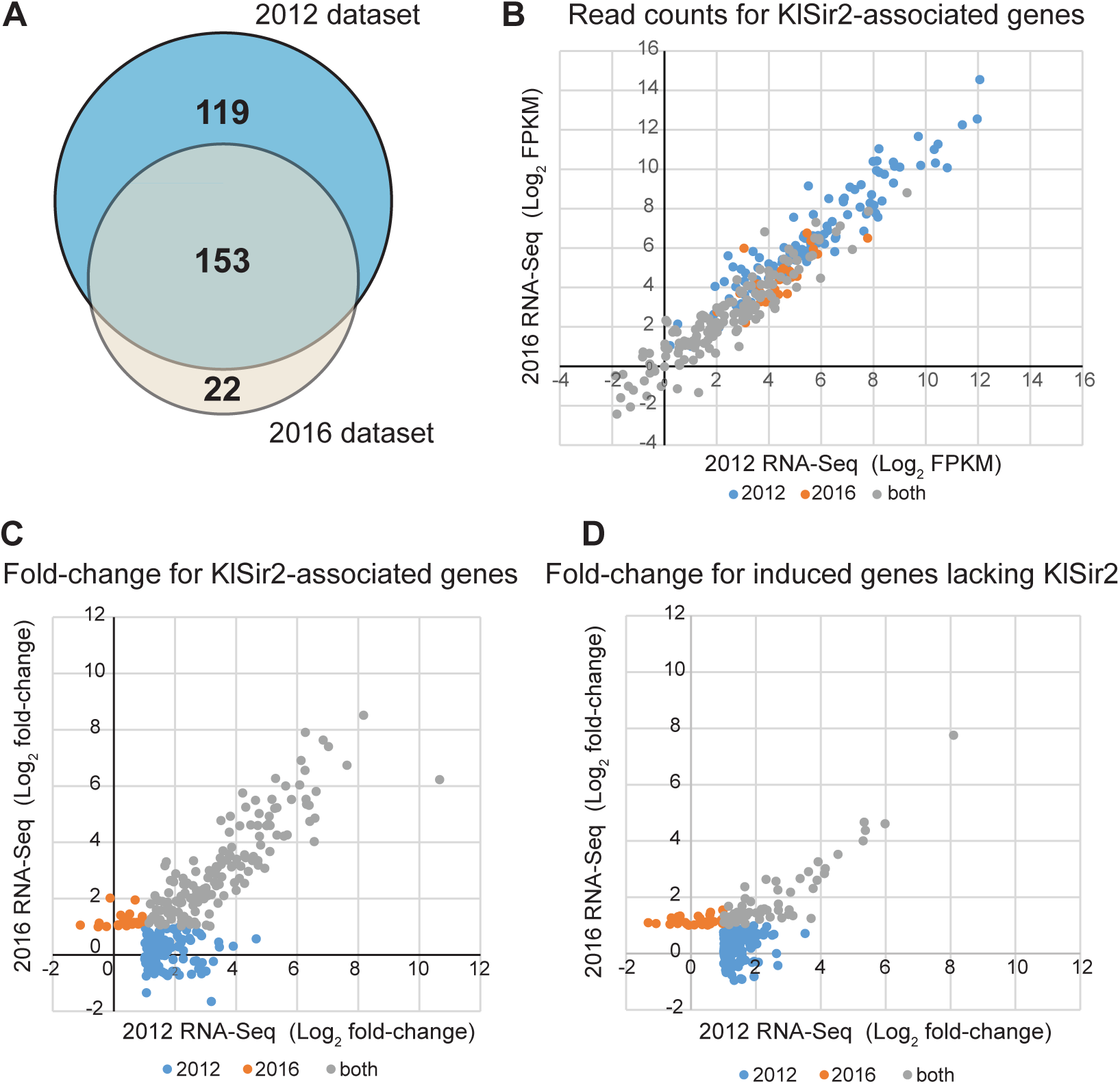
Comparison of two *K. lactis* RNA-Seq datasets. (A) The overlap was determined for KlSir2-associated genes induced in *sir2*Δ compared to wild-type cells in the 2012 RNA-Seq dataset (272) and the 2016 dataset (175). (B) For each KlSir2-associated gene induced in at least one dataset, the log_2_ FPKM (Fragments Per Kilobase of transcript per Million mapped reads) from wild-type cells from each dataset was plotted. Genes that were induced in both datasets (gray) tended to have lower expression (FPKM) than genes induced in only one of the two datasets (2012, blue or 2016, red). (C) For each KlSir2-associated gene induced in at least one dataset, the log_2_ fold-change (*sir2*Δ/wild-type) from each dataset was plotted. Genes that were induced in both datasets (gray) tended to have higher fold-change than genes induced in only one of the two datasets. (D) For induced genes that were not associated with KlSir2, the log_2_ fold-change (*sir2*Δ/wild-type) from each dataset was plotted. Fewer of these genes were induced in both datasets. These data strengthen the conclusion that the 153 KlSir2-associated genes induced in both datasets are repressed by KlSir2.

**Figure S2.**
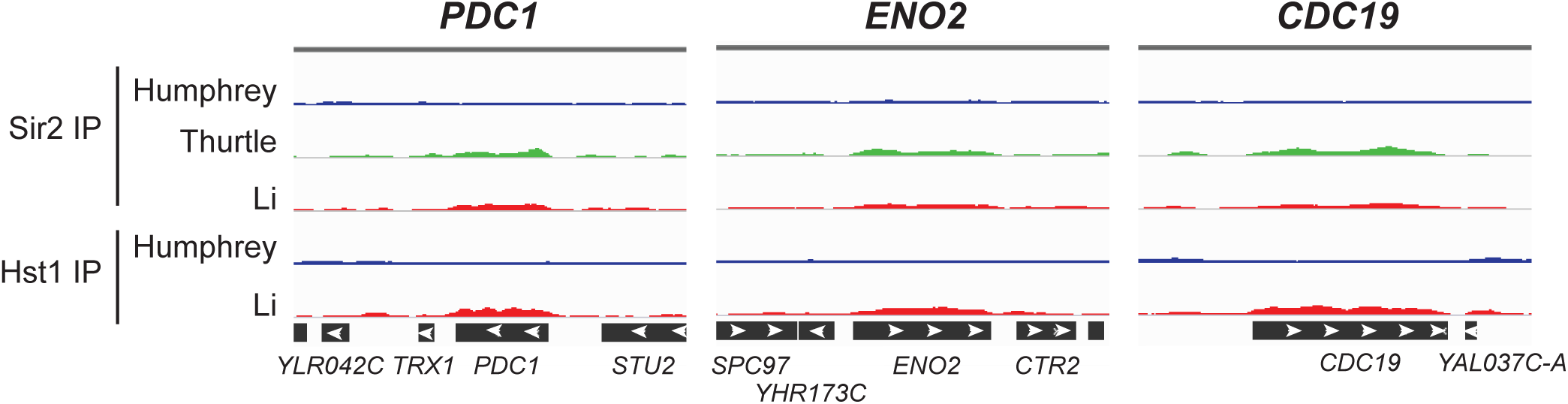
ScSir2 and ScHst1 were not enriched at highly expressed genes. ScSir2 and ScHst1 ChIP-Seq reads from this study (Humphrey) and two others (Thurtle and Rine, 2014; Li et al., 2013) were piled up at three genomic loci reported to be associated with these deacetylases (Li et al., 2013). Our data revealed little enrichment of ScSir2 or ScHst1. In the other studies, slight enrichment was observed, but could result from a known “hyper-ChIP” artifact (Teytelman et al., 2013).

**Figure S3.**
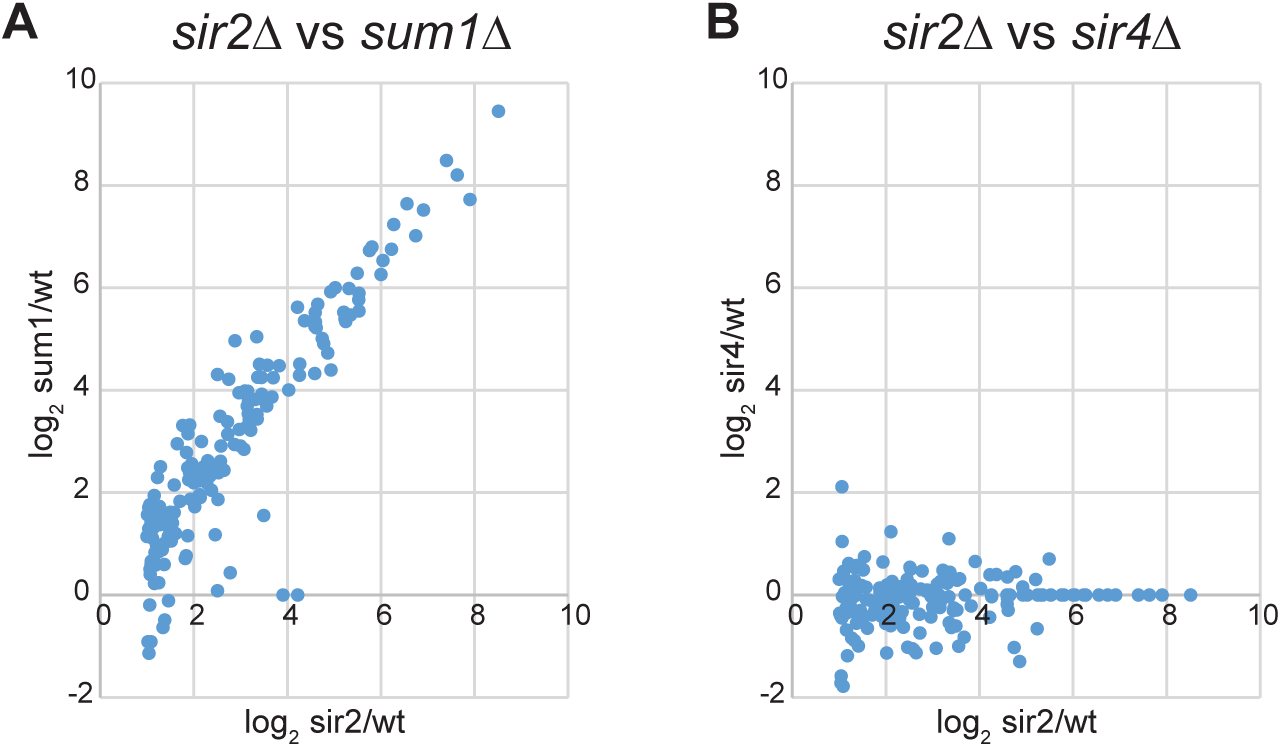
Most KlSir2-regulated genes were also regulated by KlSum1. (A) For each KlSir2-regulated gene, the log_2_ fold-change was plotted for *sir2*Δ compared to wildtype cells and *sum1*Δ compared to wildtype cells. (B) For the same genes, the log_2_ fold-change was plotted for *sir2*Δ compared to wildtype cells and *sir4*Δ compared to wildtype cells.

## SUPPLEMENTAL MATERIALS AND METHODS

### Plasmid construction

Plasmids used in this study are listed in Table S1. The *sir2Δ::NatMX* deletion cassette in pLR809 was generated by homologous recombination in yeast. Specifically, the reading frame of *KlSIR2* in pLR730 (FROYD AND RUSCHE 2011) was replaced with NatMX amplified from pAGT100 (KAUFMANN AND PHILIPPSEN 2009) using primers 5’-tagctggaactggagcgcggaatattcattatctggagttcCCAGTGAATTCGAGCTCGG and 5’-atcagatcataagtgattcaaagcaacaagatttattcaaCATGATTACGCCAAGCTTGC. For the *K. lactis* mating assay, fluorescent proteins were cloned into integrating vector pGBN19 (READ *et al*. 2007), which drives expression from the *LAC4* promoter. First, the MFα1 leader sequence was removed from pGBN19 by digesting with *Hin*DIII and *Xho*I, blunting the ends, and religating the plasmid to generate pLR1076. Next, yeast enhanced GFP (yEGFP) was excised from pKT128 (SHEFF AND THORN 2004) using *Kpn*I and *Bam*HI and ligated into the *Kpn*I and *Bgl*II sites of pLR1076 to generate pLR1087. Separately, mCherry was excised from plasmid yEpGAP-cherry-MCS (KEPPLER-ROSS *et al*. 2008) using *Hin*DIII and *Bgl*II and ligated into the *Hin*DIII and *Bgl*II sites of pLR1076 to generate pLR1085.

### Yeast strain construction

Yeast used in this study are listed in Table S2. Most *K. lactis* strains were derived from Os334 and Os335 (HEINISCH *et al*. 2010), which are congenic with the type strain CBS2359. To generate strains for RNA-Seq, we first deleted the *HM* loci. In strain Os334, *HMLα* was replaced with loxP-flanked KanMX from pCUG6 (PRIBYLOVA *et al*. 2007), and in strain Os335, *HMR**a*** was replaced with loxP-flanked *LEU2* from pJJ955L (HEINISCH *et al*. 2010). The markers were then removed by transiently transforming the yeast with pJJ958 (HEINISCH *et al*. 2010) expressing Cre recombinase and *URA3*. Next, these two strains were crossed to generate LRY2835 with both *HM* loci deleted. Finally, repressor proteins were deleted using the *sir2Δ::NatMX* deletion cassette from pLR809 to generate LRY2849 and 2850, the *sir4Δ::URA3* cassette from LRY1946 (HICKMAN AND RUSCHE 2009) to generate LRY3096, or a *sum1Δ::KanMX* cassette generated *in vitro* using the NEBuilder kit (New England Biolabs) and a KanMX cassette amplified from pFA6a-KanMX (BAHLER *et al*. 1998) to generate LRY3098. To generate prototrophic strains (LRY2992, LRY2993, LRY3027, and LRY3028), *ADE2*, *HIS3*, and *LEU2* were amplified from CK57-7A (CHEN AND CLARK-WALKER 1994) and used to transform the RNA-Seq strains as well as an isogenic *MATα* strain derived from the same cross that produced LRY2835. For the sporulation assay, diploid cells were generated by mating haploid strains that were intermediates in the construction of prototrophic strains. *MAT**a** ura3* and *MATα leu2* haploids were mated to generate diploids homozygous for the deletions of the *HM* loci. For the mating assay, the prototrophic strains were transformed with constructs to integrate fluorescent proteins under the control of the *LAC4* promoter. *LAC4*::mCherry was derived from pLR1085 cut with HpaI and XmaI, and *LAC4*::yEGFP was derived from pLR1087 cut with SacII. For the ChIP-on-chip experiment, Sir2 was tagged as previously described (HICKMAN AND RUSCHE 2009) in strain SAY538 (BARSOUM *et al*. 2010). The resulting strain was crossed to CK213-4c (KEGEL *et al*. 2006), and two of the progeny, LRY2021 and 2022, were used for chromatin IP. Ambiguities were later noted in the mating-type of LRY2022.

*S. cerevisiae* strains were derived from the standard laboratory strain W303-1b. Most were generated through transformations and crosses to recombine previously constructed alleles, including *hst1Δ::KanMX* and *sir2Δ::TRP1* (RUSCHE AND RINE 2001), *HST1::5HA-URA3* (RUSCHE AND RINE 2001; HICKMAN AND RUSCHE 2007), and *hmlαΔ::TRP1* (STONE *et al*. 2000). The *hmr**a**Δ::URA3* allele was generated by one-step gene replacement using *URA3* amplified from pRS406 with oligos 5’-GAAATGCAAGGATTGGTGATGAGATAAGATAATGAAACATagattgtactgagagtgcac and 5’-CCTCGAGGTGTAATCTAAATAATAACTTTATCGCAGTAGActgtgcggtatttcacaccg. The *SIR2::3HA-URA3* allele was generated by one-step gene insertion at the end of the *SIR2* reading frame using a 3xHA tag amplified from pLR522 (HANNER AND RUSCHE 2017) with primers 5’-ATGGAAAAAGATTTTCAAGTGAATAAGGAGATAAAACCGTAT ggcggccgcatcttttac and 5’-CAGGGTACACTTCGTTACTGGTCTTTTGTAGAATGATAAAgctcgaattcctgcagcccg.

### Yeast transformation

*S. cerevisiae* cells were transformed using the PEG-LiOAc method (SCHIESTL AND GIETZ 1989). Cells were harvested at OD_600_ around 1 and washed twice with 0.1 volumes of TEL (10 mM tris, pH 7.5, 1 mM EDTA, 100 mM LiOAc). Cells were resuspended in TEL at 10 μl/OD cells, and 100 μl of cells were added to 0.1 μg of linear DNA plus 30 μg sheared salmon sperm DNA. Cells were incubated at 30° for 30 minutes, combined with 750 μl 40% PEG-TEL, and incubated at 30° for 30 minutes. Finally, cells were heat shocked at 42° for 10 minutes and plated on selective medium. *K. lactis* cells were transformed using electroporation (HICKMAN AND RUSCHE 2009). Briefly, cells were harvested at an optical density around 1 and resuspended at 15 OD/ml in YPD containing 25 mM DTT and 20mM HEPES, pH 8. Cells were shaken for 30 minutes at 30°, collected, and washed in electroporation buffer (10mM tris pH7.5, 270mM sucrose, 1mM LiOAc). Cells were then resuspended at 100 OD/ml in electroporation buffer. Electroporation reactions were set up in 0.2 mm cuvettes using 50 μL cells, 1 μL 10 mg/mL salmon sperm DNA, and 0.5-1 μg DNA in a volume no more than 5 μL. Electroporation conditions were 1000 V, 300 Ω, and 25 μF. After electroporation, cells were incubated in YPD four hours at 30° and then spread on selective media.

### Chromatin IP and processing for microarray or sequencing

For the ChIP on Chip experiment from *K. lactis*, chromatin IP was conducted as previously described (HICKMAN AND RUSCHE 2009), with some exceptions. Cells were crosslinked for one hour each in 10 mM DMA and then 1% formaldehyde. After cell lysis, chromatin was sonicated four times for 15 seconds. 160 μl of lysate (derived from 10 OD equivalents of cells) was brought to a final volume of 400 μl in lysis buffer and incubated overnight with 7 μl anti-HA antibody (Upstate). The immunoprecipitated DNA was labeled with either Cy5-or Cy3-conjugated dUTP (Perkin Elmer NEL578001EA or NEL579001EA), using Klenow DNA polymerase (NEB M0212M) and random nonamer oligonucleotides (IDT). 500 ng of input DNA or an entire immuno-precipitated DNA sample was dried in a speed-vac and resuspended in 15 μL of primer mix (1X NEB buffer 2, 5 μg of random nonamer). Once the DNA was dissolved, 2 nmole of labeled dUTP was added in 2 μl. The samples were placed in a thermocycler and denatured for 5 minutes at 95°, and then cooled to 4°. The samples were combined with 3 μL of Klenow reaction mix, resulting in a final concentration of 1X NEB buffer 2, 0.25 mM dATP, 0.25 mM dCTP, 0.25 mM dGTP, 0.1 mM dTTP and 12.5 U of Klenow. The sample was ramped to 37° at 0.1°/sec and then incubated for 30 minutes. Following incubation, the sample was heat-denatured and cooled to 4°, and fresh Klenow (4 U) was added for a second round of labeling. Finally, unincorporated nucleotides, oligonucleotides, and dye were removed using Microcon YM-30 filters (Millipore). Labeled DNA was hybridized to the tiled Agilent array in hybridization buffer overnight at 65°. The microarray was washed and scanned according the manufacturer’s instructions.

For the ChIP-Seq experiment from *S. cerevisiae*, chromatin IP was performed essentially as described (RUSCHE AND RINE 2001). Cells were harvested at OD_600_ around 1. Cells were crosslinked for one hour each in 10 mM DMA and then 1% formaldehyde. The immunoprecipitation was conducted with 10μL of Protein A agarose beads in the absences of BSA and salmon sperm DNA. Library preparation and sample barcoding was done at the Next-Generation Sequencing facility at University at Buffalo. The samples were then sequenced on an Illumina HiSeq2500 using 50 bp single-end sequencing.

## LIST OF TABLES

Tables S1 and S2 are included in this document. Tables S3-S9 are posted on FigShare.

**Table S1.**
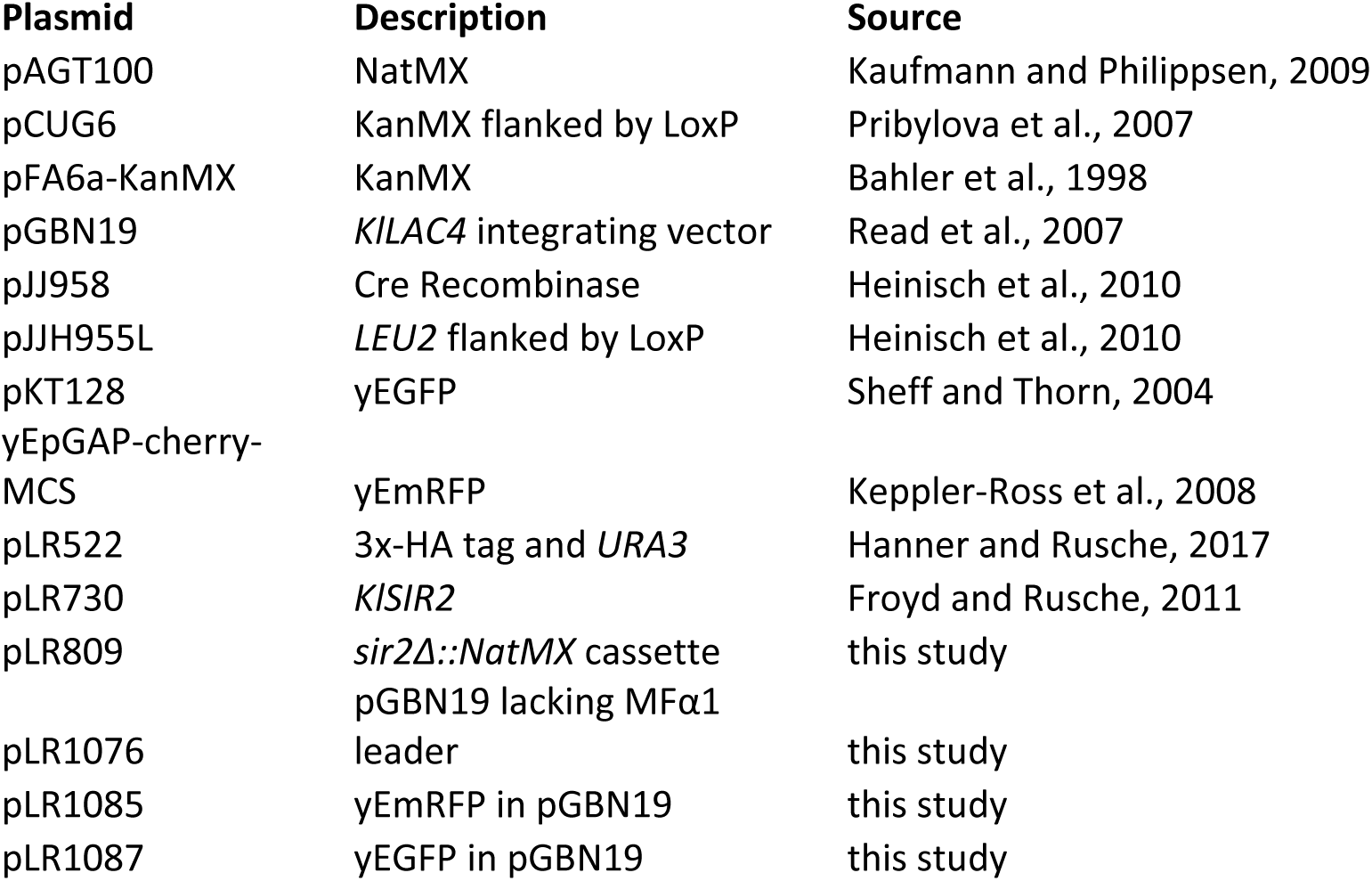
Plasmids used in this study

**Table S2.**
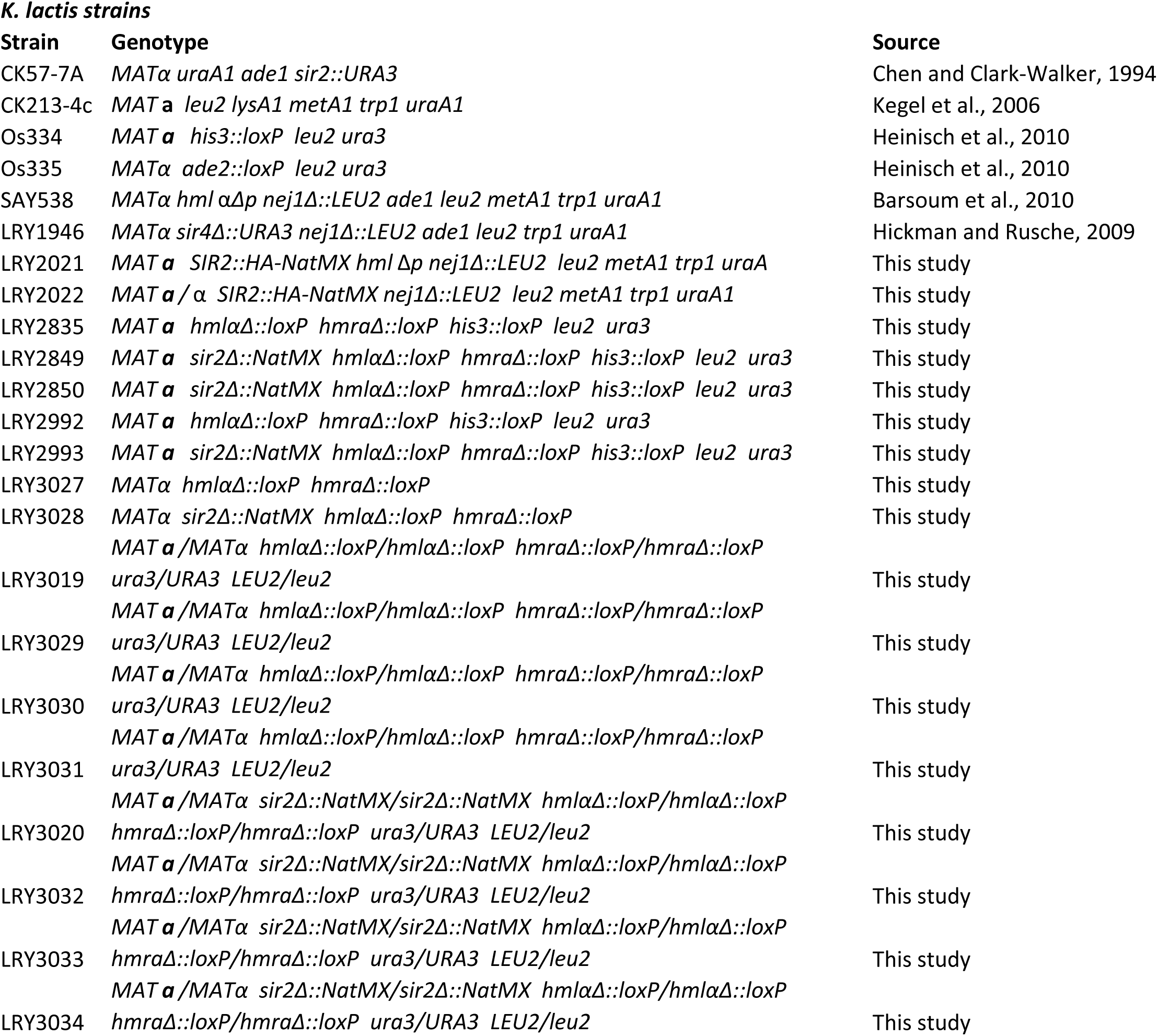
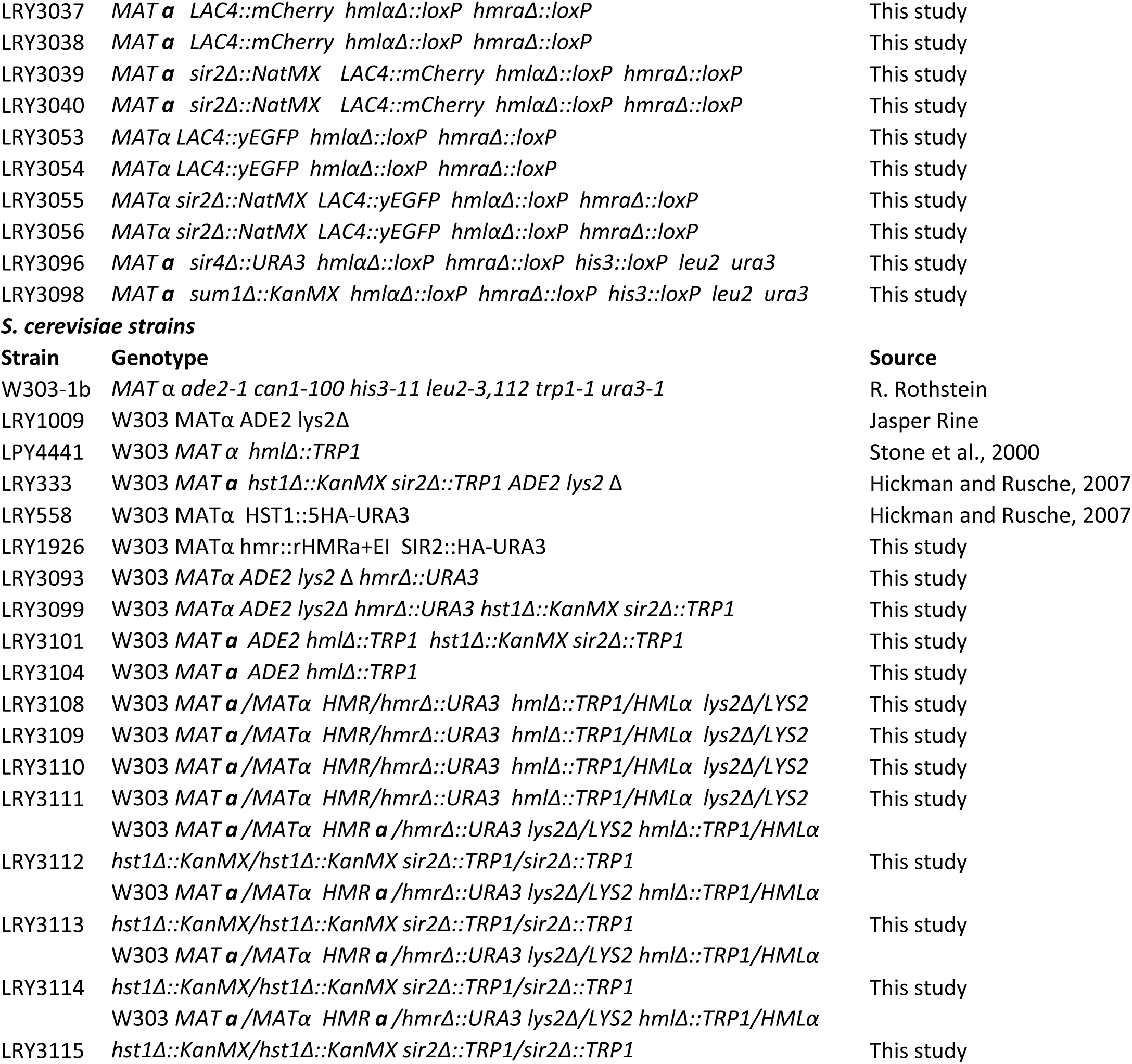
Yeast strains used in this study

**Table S3. RNA-Seq and ChIP-Seq data for all *S. cerevisiae* genes**

Raw data for all annotated *S. cerevisiae* genes.

**Table S4. RNA-Seq and ChIP-Chip data for all *K. lactis* genes**

Raw data for all annotated *K. lactis* genes.

**Table S5. ScSir2-regulated genes**

Each row represents a gene that was both associated with ScSir2 and upregulated at least two-fold in *sir2Δ hst1Δ* compared to wild-type *S. cerevisiae*. The KlSir2-regulated column indicates whether the *K. lactis* ortholog is regulated by KlSir2 based on both RNA-Seq datasets (12&16) or just the newer dataset (2016). The Ellahi column indicates whether the gene was identified by (ELLAHI *et al*. 2015) as SIR-regulated.

**Table S6. ScHst1-regulated genes**

Each row represents a gene that was both associated with ScHst1 and upregulated at least two-fold in *sir2Δ hst1Δ* compared to wild-type *S. cerevisiae*. The KlSir2-regulated column indicates whether the *K. lactis* ortholog is regulated by KlSir2 based on both RNA-Seq datasets (12&16) or just the older dataset (2012). The Bedalov column indicates whether the gene was identified by (BEDALOV *et al*. 2003) as Hst1-regulated. The McCord columns indicate whether the gene was identified by (MCCORD *et al*. 2003) as Hst1-or Sum1-regulated. The Borde and Friedlander columns indicate whether the gene was increased during sporulation in two expression studies (FRIEDLANDER *et al*. 2006; BORDE *et al*. 2009). The categories and subcategories were developed manually based on GO terms and functional information about each gene.

**Table S7. KlSir2-regulated genes identified using 2016 RNA-Seq data**

Each row represents a gene that was both associated with KlSir2 and upregulated in the 2016 dataset at least two-fold in *sir2Δ* compared to wild-type *K. lactis*. The *S. cerevisiae* orthologs were determined through a reciprocal BLASTP procedure followed by manual refinement, as described in the methods. For genes whose top *S. cerevisiae* BLASTP hit was more similar to another *K. lactis* gene, no *S. cerevisiae* ortholog is given. Instead, the description indicates that the gene is related to its top hit. The 2012 column indicates whether the gene was also induced in the 2012 RNA-Seq dataset. The categories and subcategories were developed manually based on GO terms and functional information about each gene and its *S. cerevisiae* ortholog.

**Table S8. KlSir2-regulated genes identified using 2012 RNA-Seq data**

Each row represents a gene that was both associated with KlSir2 and upregulated in the 2012 dataset at least two-fold in *sir2Δ* compared to wild-type *K. lactis*. Genes that were also upregulated in the 2016 dataset are excluded from this list and can be found in Table S7. Columns are as described for Table S7.

**Table S9. RNA-Seq and ChIP-Seq data for metabolic genes**

This table is the basis for Figure 3 and includes all *S. cerevisiae* genes known to act in the each pathway included in the figure. For each gene and its *K. lactis* ortholog, data are provided for the association with ScSir2, ScHst1, and KlSir2 and the expression change in deletion compared to wild-type cells.

